# The tree labeling polytope: a unified approach to ancestral reconstruction problems

**DOI:** 10.1101/2025.02.14.638328

**Authors:** Henri Schmidt, Benjamin J. Raphael

## Abstract

**Motivation:** Reconstructing unobserved ancestral states of a phylogenetic tree provides insight into the history of evolving systems and is one of the fundamental problems in phylogenetics. For a fixed phylogenetic tree, the most parsimonious ancestral reconstruction – a solution to the small parsimony problem – can be efficiently found using the dynamic programming algorithms of Fitch-Hartigan and Sankoff. Ancestral reconstruction is important in many applications including inferring the routes of metastases in cancer, deriving the transmission history of viruses, determining the direction of cellular differentiation in organismal development, and detecting recombination and horizontal gene transfer in phylogenetic networks. However, most of these applications impose additional *global* constraints on the reconstructed ancestral states, which break the local structure required in the recurrences of Fitch-Hartigan and Sankoff.

**Results:** We introduce an alternative, polyhedral approach to ancestral reconstruction problems using the *tree labeling polytope*, a geometric object whose vertices represent the feasible ancestral labelings of a tree. This framework yields a polynomial-time linear programming algorithm for the *small parsimony problem*. More importantly, the tree labeling polytope facilitates the incorporation of additional constraints that arise in modern ancestral reconstruction problems. We demonstrate the utility of our approach by deriving mixed-integer programming algorithms with a small number of integer variables and strong linear relaxations for three such problems: the parsimonious migration history problem, the softwired small parsimony problem on phylogenetic networks, and the convex recoloring problem on trees. Our algorithms outperform existing state-of-the-art methods on both simulated and real datasets. For instance, our algorithm scales to trace routes of cancer metastases in trees with thousands of leaves, enabling the analysis of large trees generated by recent single-cell sequencing technologies. On a mouse model of metastatic lung adenocarcinoma, the tree labeling polytope allows us to infer simpler migration histories compared to previous results.

**Availability:** Python implementations of the algorithms provided in this work are available at: github.com/raphael-group/tree-labeling-polytope.

## 1 Introduction

A foundational problem in phylogenetics is to infer the ancestral states of phylogenetic characters from a phylogenetic tree; i.e. given a leaf-labeled tree to label the ancestral vertices in the tree in order to optimize some objective. Finding the most parsimonious labeling is called the *small parsimony problem* and is solved in polynomial time via the Fitch-Hartigan recurrence [1, 2] in the unweighted case, and Sankoff’s algorithm [3–5] in the weighted case. Classically, this inference aims to reconstruct the molecular sequences of ancestral species [6–8]. Over the past decade, however, the importance of ancestral reconstruction has grown beyond its original scope, finding numerous applications across subfields of biology. For example, the perspective of ancestral reconstruction has been successfully applied to infer the metastatic history of tumors [9–15], host-to-host transmission networks of viruses [16–25], and cellular differentiation maps in developmental biology [26]. In modeling recombination and horizontal gene transfer, ancestral reconstruction has been used to score and rank different phylogenetic network hypotheses [27–30]. In inferring evolutionary trees, ancestral reconstruction is used to score and order different trees within a parsimony framework, providing the basis of hundreds of algorithms for inferring phylogenetic trees.

Sankoff’s algorithm [3–5] efficiently finds the most parsimonious ancestral labeling when no constraints are imposed on the inferred labeling. However, many *parsimonious* ancestral reconstruction problems of interest [11, 19, 20, 22, 24, 27–30] introduce *global constraints* or *additional criteria*, yielding NP-hard variants of the classic small parsimony problem. As a result, naïvely applying Sankoff’s algorithm results in brute-force approaches with exponential runtime, making it impractical for large datasets. To address this, researchers have developed specialized methods, including integer linear programming approaches [11, 26, 31–34] and dynamic programming algorithms [24, 27, 30, 35–37], though both methods come with their own drawbacks.

In the combinatorial optimization literature, however, there is an alternative approach which starts with a linear programming formulation of the unconstrained problem and tacks on the additional, typically integer, complicating constraints. For example, the minimum spanning tree problem has an elegant linear programming formulation solvable in polynomial time, of which the feasible region is the *spanning tree polytope* due to Edmonds [38, 39]. However, when constraints on the degree of the inferred spanning tree are added, the minimum spanning tree problem becomes NP-hard [40]. To solve the NP-hard bounded degree minimum spanning tree problem, state-of-the-art algorithms [40, 41] start by simply appending the degree constraints to the linear program for the (unconstrained) minimum spanning tree problem. Solving the resultant linear program, one obtains non-integral solutions, which are then fixed using either rounding [40–42] or cutting plane [43, 44] approaches.

The advantage of building an integer programming formulation by starting with a linear program for the easy subproblem is that one obtains formulations with strong linear relaxations which often contain only a small number of integer variables [45, 46]. In fact, integer programming formulations constructed in this way serve as the starting point for state-of-the-art approximation and exact algorithms for solving NP-complete generalizations of minimum spanning tree [40, 47, 48], shortest path [49, 50], and maximum flow problems [51–53]. Unfortunately, it is often not obvious how to obtain linear programming formulations, or *polyhedral descriptions*^1^, of combinatorial problems.

### Contributions

As many parsimonious ancestral reconstruction problems of interest are variants of the small parsimony problem with additional, complicating constraints, we sought to develop a polynomial sized, polyhedral description of the small parsimony problem. To this end, we construct a polyhedral description of the set of ancestral labelings over a fixed tree by defining a new geometric object called the *tree labeling polytope* (Figure 1a). By studying the basic properties of the tree labeling polytope, we show that it is described by a polynomial-sized set of 𝒪 (*nm*^2^) linear inequalities (or *facets*), where *m* is the alphabet size and *n* is the size of the phylogenetic tree. To obtain this result, we provide an algorithmic primal-dual packing proof of the total dual integrality of a corresponding *tree labeling system*. As an immediate corollary, we obtain, to the best of our knowledge, the first polynomial-time algorithm for the small parsimony problem based on linear programming.

**Figure 1:**
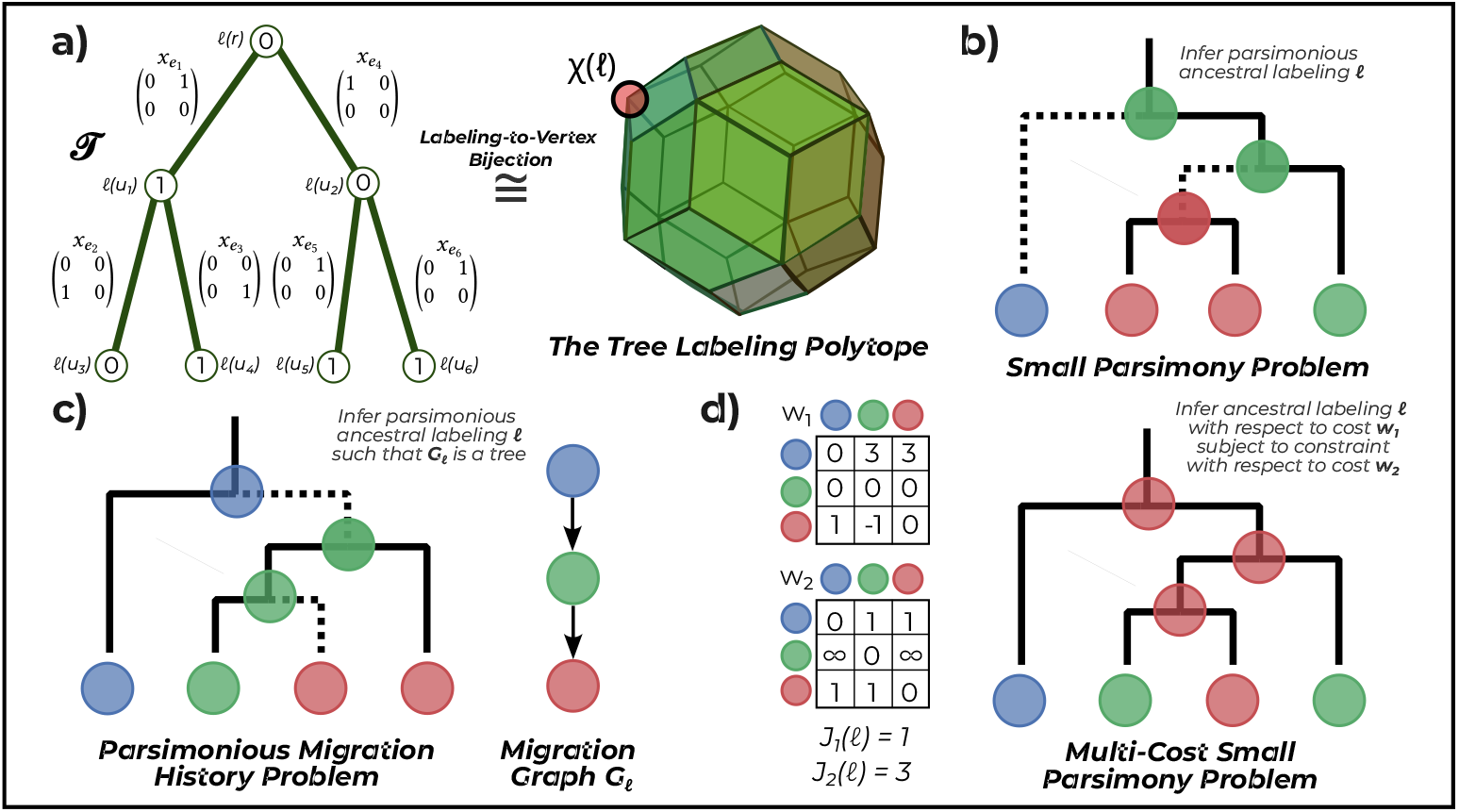
**a)** The vertices of the tree labeling polytope 𝒫 (𝒯)denoted as χ(*ℓ*) are in one-to-one correspondence with the vertex labelings of *l* of 𝒯. **b**) The small parsimony problem aims to find the ancestral labeling with the fewest number of label changes. **c)** The parsimonious migration history problem aims to find the most parsimonious ancestral labeling under structural constraints on the induced migration graph *G*_*ℓ*_. **d)** The multi-cost small parsimony problem finds the cheapest labeling with respect to one cost function *w*_1_ while satisfying a constraint on a second cost function *w*_2_.

We demonstrate the utility of the tree labeling polytope by developing mixed integer linear programming algorithms to solve three generalizations of the small parsimony problem, each of which is NP-complete:

- We solve a generalization of the *parsimonious migration history problem* introduced in [11] which introduced the MACHINA algorithm. In particular, we develop a fixed parameter tractable, mixed integer linear programming formulation for this generalization with *strong* linear relaxations.
- We solve the softwired small parsimony problem on phylogenetic networks [27, 28, 30] with a fixed parameter tractable, mixed integer linear programming formulation containing only *r* binary variables, where *r* is the number of *reticulation*, or in-degree 2, vertices in the network.
- We solve an analog of the convex recoloring problem on trees [54] called the *multi-cost small parsimony problem*. By drawing a connection to the *constrained minimum spanning tree* approximation algorithm of Ravi and Goemans [47], we obtain a mixed integer programming formulation for the multi-cost small parsimony problem with strong linear programming relaxations, which we extend to the convex recoloring problem.

We demonstrate the practical efficacy of our approach by developing scalable algorithms for all three of the afore-mentioned problems which outperform state-of-the-art algorithms on both simulated and real datasets. For the parsimonious migration history problem, our algorithm scales to trees with thousands of leaves, making it possible to analyze large trees obtained from single-cell sequencing technologies, compared to existing algorithms which scale to phylogenetic trees with a hundred leaves [11]. Next, we demonstrate the utility of the tree labeling polytope for generating biological insights by inferring simpler migration histories than those previously reported in 83 tumor clones derived from single-cell lineage tracing data from a mouse model of lung adenocarcinoma [15]. Further, using a precise quantitative model based on convex recolorings, we find that while some tumors suggest patterns of monoclonal seeding, most do not, concordant with recent literature on the polyclonality of metastatic tumors [55, 56].

## 2 Parsimonious ancestral labeling problems

Let 𝒯 = (*V* (𝒯), *E* (𝒯)) be a rooted tree with root *r* = *r* (𝒯), leaf set *L*(𝒯), and *n* edges. For a vertex *u ∈ V* (𝒯), let *δ* (*u*) denote the set of children of *u*, Δ(*u*) be the set of descendants of *u*, and *ρ* (*u*) be the parent of vertex *u* in *V* (𝒯) (*ρ* (*r*) is undefined).

We are interested in describing labelings of the vertices of 𝒯. Assume Ω = {1, …, *m*} is an alphabet of *m* labels. A map *ℓ* : *L*(𝒯) ⟶ Ω is a *leaf labeling* of 𝒯 ^2^. A map 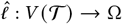 is a *vertex labeling* of 𝒯 and 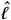 is an *extension* of the leaf labeling *ℓ* if 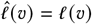 for all leaves *𝓋 ∈ L* (𝒯). A *cost function* is a map *w* : *E* (𝒯) × Ω × Ω → ℝ ∪ {∞}, and gives the cost of a transition from label *i* to *j* across edge *e*. A cost function *w* is *integral* if range (*w*) ⊆ ℤ and *non-integral* otherwise. A cost function *w* is called *edge independent* if *w* (*e, i, j*) = *w* (*e*^′^, *i, j*) for all pairs of edges *e, e*^′^ ∈ *E* (𝒯), and is called *edge dependent* otherwise. A cost function *w* is *normal* if *w* (*e, i, i*) = 0 for all edges *e E* and *not normal* otherwise. A normal cost function *w* is called *label independent* if *w* (*e, i*_1_, *j*_1_) = *w* (*e, i*_2_, *j*_2_) for all *i*_1_ ≠ *j*_1_ and *i*_2_ ≠ *j*_2_, and is called *label dependent* otherwise.

### 2.1 Problem statements

The small parsimony problem [57] asks to find an extension 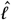 of the leaf labeling *ℓ* such that the total weighted cost is minimized. In other words, the small parsimony problem finds the *most parsimonious* ancestral explanation of the leaves (Figure 1b).

#### Problem

(Small Parsimony). *Given a tree* 𝒯, *a label set* Ω, *a leaf labeling ℓ, and a cost function w* : Ω × Ω → ℝ, *find an extension* 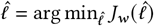 *where* 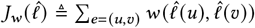.

Originally, the unweighted small parsimony problem (taking *w*(*i, j*) = 𝟙 (*i* ≠ *j*) was solved in 𝒪 (*nm*) time by the famous Fitch-Hartigan algorithm [1, 2]. More generally, Sankoff gave a 𝒪 (*nm*^2^) time algorithm for the case where the cost of a transition from label *i* to label *j* can depend on the labels [3, 4]. In fact, essentially all dynamic programming algorithms for different cases of the small parsimony problem can be viewed as specializations, and often optimizations, of the *Sankoff-Rousseau recurrence* [3, 4], an idea fleshed out in detail in [5].

Generalizing slightly, Sankoff’s algorithm need not require that the inferred vertex labeling 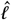 is an extension of *ℓ*, nor that the cost function is constant with respect to the edge. In particular, Sankoff’s algorithm also solves the version where the cost function is a map *w* : *E* (𝒯) × Ω × Ω → ℝ ∪ {∞} and the cost 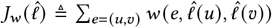. To recover the case where 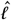 is required to extend *ℓ*, simply set the cost function *w* (*e, i, j*) = ∞ if *e* = (*u, 𝓋*) is a pendant edge with *j* ≠ *ℓ* (*𝓋*). Throughout the remainder of the manuscript, all references to the small parsimony problem refer to this more general setting.

In several applications, additional constraints [11, 16, 20–22, 24, 25] are placed on the inferred labeling. For example, [11] places structural constraints on the inferred labeling 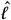, defining the *parsimonious migration history problem* (Figure 1c). In this setting, the label set Ω is interpreted as a set of *m anatomical sites*, the vertices of 𝒯 are interpreted as *cells*, and the labeling *ℓ* (*𝓋*) is interpreted as the *anatomical location* of cell *𝓋*. The main quantity of interest is the *migration graph* 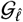 associated with a vertex labeling 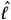, which tells us the patterns of cellular migration between anatomical sites. The *migration graph* 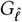 has vertex set Ω and includes a directed edge (*i, j*) if and only if there exists an edge (*u, 𝓋*) ∈ *E* (𝒯)such that 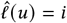 and 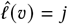. The parsimonious migration history problem aims to find the most parsimonious ancestral labeling 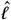 under structural constraints on the induced migration graph 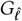, specified with a graph class 𝒢 (Figure 1c).

#### Problem

(Parsimonious Migration History). *Given a tree 𝒯, a label set* Ω, *and a cost function w, find the vertex labeling* 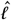 *with minimum cost* 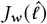 *such that* 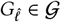.

El-Kebir et al. [11] considered several choices of the graph class 𝒢 including the class of rooted trees, the class of directed acyclic graphs, and the class of all graphs^3^. When 𝒢 consists of all graphs, the parsimonious migration history problem is equivalent to the small parsimony problem and thus solveable in polynomial time. Otherwise, the problem is more difficult. For example, the parsimonious migration history problem is NP-hard when 𝒢 is the class of trees [35].

Another variant of the small parsimony problem is the *softwired small parsimony problem* [27, 28], which generalizes the small parsimony problem to phylogenetic networks, modeling events such as horizontal gene transfer and recombination [58]. A *phylogenetic network* 𝒩 = (*V* (𝒩), *E* (𝒩)) is a directed acyclic graph with a unique root *r* (𝒩) having in-degree zero and a leaf set *L*(𝒩). We assume that each vertex *𝓋* in 𝒩 has in-degree at most two, and use *R* (*V*) to denote the set of vertices with in-degree two, herein referred to as *reticulation vertices*. The set *R* (*E*) of edges with terminals in *R* (*V*) are the *reticulation edges*. For a spanning tree 𝒯 of 𝒩, let 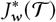 denote the small parsimony score of 𝒯. Using this notation, the softwired small parsimony problem is defined as follows.

#### Problem

(Softwired Small Parsimony). *Given a phylogenetic network* 𝒩 = (*V, E*), *a label set* Ω, *and a cost function w, compute the* softwired small parsimony score:

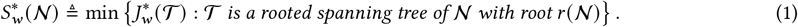

That is, the softwired small parsimony score is the minimum parsimony score over all spanning trees of 𝒩. As each spanning tree of 𝒩 corresponds to the choice of one of two in-edges of *𝓋* for all reticulation vertices *𝓋 ∈ R* (*V*), the total number of spanning trees is 2^|*R* (*V*) |^. In the case when 𝒩 is a tree, 𝒩 contains only a single spanning tree (itself) and the softwired small parsimony problem reduces to the small parsimony problem. The softwired small parsimony problem is NP-complete and not-approximable, meaning that no polynomial time algorithm admits a constant factor approximation assuming *P* ≠ *N P* [31]. Numerous algorithmic approaches have been developed for the softwired small parsimony problem including specialized dynamic programming [27, 28, 30, 59], integer programming [31, 37], and constant-factor approximation algorithms [37].

Finally, we introduce the *multi-cost small parsimony problem*, a cost constrained variant of the small parsimony problem, which to our knowledge is not yet studied in the phylogenetics literature (Figure 1d). In this setting, we are provided two cost functions *w*_1_, *w*_2_ and a real number *k*, and the goal is to minimize the primary cost 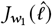 subject to the constraint 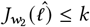 *k* on the secondary cost. This problem is analogous to the constrained minimum spanning tree [40, 47] problems studied in the operations research and computer science literature, which has an additional constraint on the spanning tree with respect to a second set of weights. While the minimum spanning tree problems is solved in polynomial time with dynamic programming algorithms, the constrained minimum spanning tree problem is NP-complete [60]. Similarly, the multi-cost small parsimony problem is NP-complete. This problem is also closely related to the NP-complete convex recoloring problem [54], an idea we describe in (Section A.3.4).

## 3 The tree labeling polytope

In this section, we describe the *tree labeling polytope*, a polytope whose vertices correspond to labelings of 𝒯. We prove that the tree labeling polytope is described by a small set of 𝒪(*nm*^2^) linear inequalities, which we call the *tree labeling system*. We defer proofs of our main theorems to Supplementary Results A due to space constraints.

### 3.1 Preliminaries

We introduce basic notation, definitions, and results from polyhedral theory, using the notation of Schrijver [61]. A vector is *integral* (resp. *rational*) if *x ∈* ℤ^*n*^ (resp. *x ∈* ℚ^*n*^). Similarly, a matrix is *integral* (resp. *rational*) if *A ∈* ℤ^*m*×*n*^ (resp. *A ∈* ℚ^*m*×*n*^). A *polyhedron* is a set of the form {*x* : *Ax* ≤ *b*}, where *A* is a matrix and *b* is a vector. A polyhedron *P* is *integral* if every face of *P* contains an integral vector. A *polytope* is the convex hull of a finite set of vectors. Equivalently, a polytope is a bounded polyhedron. The *convex hull* of a finite set of vectors *S* is denoted by conv (*S*). A vector *x* is an *extreme point* of a polyhedron *P* if it cannot be written as a convex combination of any distinct set of points in *P*. A linear system *Ax* = *b, x ≥* 0 is *totally dual integral* if for every integral vector *c*, any finite solution to max{*b*^*T*^*y* : *y*^*T*^ *A* ≤ *c*^*T*^} is attained by an integral vector.

#### Proposition 1

([61]). *Let Ax* = *b, x ≥* 0 *be a rational linear system that is totally dual integral. Then, the polyhedron P* = {*x* : *Ax* = *b, x ≥* 0} *is integral*.

### 3.2 Results

To write the small parsimony problem as a linear program, the naïve starting point is to directly encode the extensions 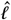 as binary vectors on the vertices of 𝒯. Specifically, given a labeling 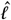 define an [(*n* + 1) *m*]-dimensional binary vector 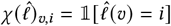 which records the label of each vertex. Unfortunately, with this encoding the cost 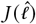 of the labeling is not a *linear function* of the vector 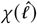. Consequently, this encoding does not allow us to write the small parsimony problem as a linear program without additional variables^4^.

Rather than encode the labeling on the vertices of 𝒯, we encode the labelings 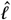 *on the edges* of 𝒯. That is, we encode 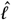 as an *nm*^2^-dimensional binary vector 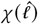 defined as 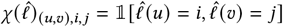. Stated simply, this encoding records the change in label (or lack thereof) across each edge. While this encoding requires 𝒪(*nm*^2^) binary variables compared to the 𝒪(*nm*) variables for the vertex encoding, the advantage is that the cost of a labeling 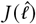 is a linear function of an *nm*^2^ -dimensional cost vector. In particular, we have that

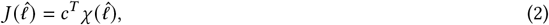

where the vector *c* is defined as *c*_*e,i,j*_ = *w* (*e, i, j*). The observation that the cost of a labeling 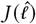 is a linear function of the indicator vector 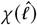 leads us to our definition of the *tree labeling polytope*.

#### Definition 1.

*Let* 𝒯 *be a rooted tree. The* tree labeling polytope 𝒫(𝒯) *is the convex hull of* 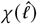, *where* 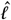 *ranges over all labelings of* 𝒯. *That is*,

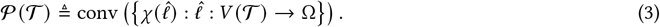

Using this notation, the (generalized) small parsimony problem is equivalent to the following linear optimization problem:

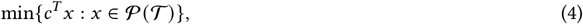

where the vector *c* is defined as *c*_*e,i,j*_ = *w* (*e, i, j*). This follows from (2) and the fact that the optimum of a linear program is always at an extreme point of its feasible region.

A final ingredient in deriving a polynomial-sized linear program is to describe the tree labeling polytope 𝒫 (𝒯)with a polynomial number of linear inequalities. From a computational perspective, the description as the convex hull of the encoded vertex labelings given in (3) is intractable as it requires enumerating all *m*^*n*+1^ vertex labelings. Instead, we aim to describe the tree labeling polytope using a polynomial (in *n* and *m*) number of linear inequalities, as this would enable us to solve the small parsimony problem in polynomial time (in *n* and *m*) using linear programming.

Our starting point is a set of necessary linear inequalities satisfied by all vectors *x ∈* 𝒫(𝒯). Namely, we define the following system of 𝒪(*nm*^2^) linear inequalities satisfied by all vectors in the tree labeling polytope.

#### Definition 2.

*Let* 𝒯 *be a rooted tree and* Ω = {1, …, *m*} *be a label set. The* tree labeling system *corresponding to is* 𝒯 *the following system of linear inequalities:*

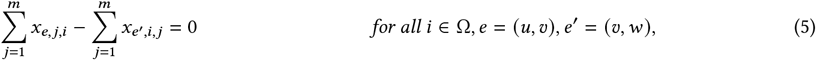

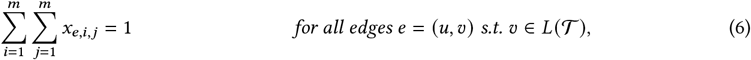

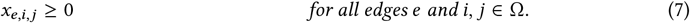

*We denote the system of linear equalities* (5)*-*(6) *by Ax* = *b and call A the* tree labeling matrix.

The intuition behind the above set of inequalities is the following. In the case where *x*_*e,i,j*_*∈* {0, 1}, equation (5) requires that each (non-root) vertex has only a single label, while equations (6)-(7) enforce that *x*_*e,i,j*_ = 1 for exactly one pair *i, j ∈* Ω. This set of necessary linear inequalities exactly describes the tree labeling polytope, in the sense of the following two results.

#### Theorem 1.

*Given a rooted tree* 𝒯 *and a label set* Ω, *the corresponding tree labeling system Ax* = *b, x ≥* 0 *is totally dual integral*.

To formally state the second result, define the augmented tree 𝒯 ^′^ of 𝒯 as the rooted tree obtained by appending the root *r* ^′^ and the edge (*r* ^′^, *r*) to 𝒯. For a subset *F* ⊆ *E* of edges, let *x*_*F*_ be the subvector of *x* consisting only of entries *x*_*f, i,j*_ with *f ∈ F*. The following result then states that the tree labeling polytope 𝒫(𝒯) is the projection of the feasible region of the tree labeling system for the augmented tree 𝒯 ^′^.

#### Theorem 2.

*Given a rooted tree* 𝒯 *and a label set* Ω, *let Ax* = *b, x ≥* 0 *be the tree labeling system corresponding to*𝒯 ^′^. *Then, the tree labeling polytope is*

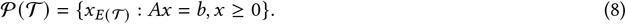

The reason for using an extended formulation [62] (i.e. defining 𝒫(𝒯) as the projection of a higher-dimensional polytope) in the preceding theorem is to ensure that the root vertex *r* has only a single label while keeping the formulation of size 𝒪(*nm*^2^). In particular, while enforcing the root vertex to have a single label in the original space is possible, it would require 𝒪(*n*^2^*m*^2^) additional constraints, increasing the size of the description by a factor of *n*.

## 4 Algorithms

Using the tree labeling polytope we derive a polynomial time linear programming algorithm for the small parsimony problem – to our knowledge, the first such algorithm of its kind. More importantly, however, using the tree labeling polytope we derive *good* mixed integer linear programming formulations of several constrained parsimony problems.

Classically, a mixed-integer programming formulation is *easy to solve* or *good* if it has *i)* a small number of integer variables and *ii)* a strong linear relaxation [46]. The linear relaxation of a mixed integer program *P* is *stronger* than that of a mixed integer program *Q* if (for a minimization problem) the objective value of the linear relaxation of *P* is greater than or equal to that of *Q*. The reason for desiring *i)* and *ii)* is because state-of-the-art solvers [64] for mixed integer programming rely on two main techniques: cutting planes, dating back to Dantzig [65], and branch-and-bound, originally due to Markowitz [66]. The effectiveness of both techniques, however, depends intimately upon *i)* and *ii)*. Specifically, strong mixed integer programs with only a small number of integer variables require fewer cutting planes and fewer branches, making the formulation easier to solve.

It is in this preceding sense we obtain *good* mixed-inter programming formulations for the parsimonious migration history, softwired small parsimony, multi-cost small parsimony, and convex recoloring problems using the tree labeling polytope. Due to space constraints, these results are described in Supplementary Results A.3 and summarized in Table 1.

**Table 1:**
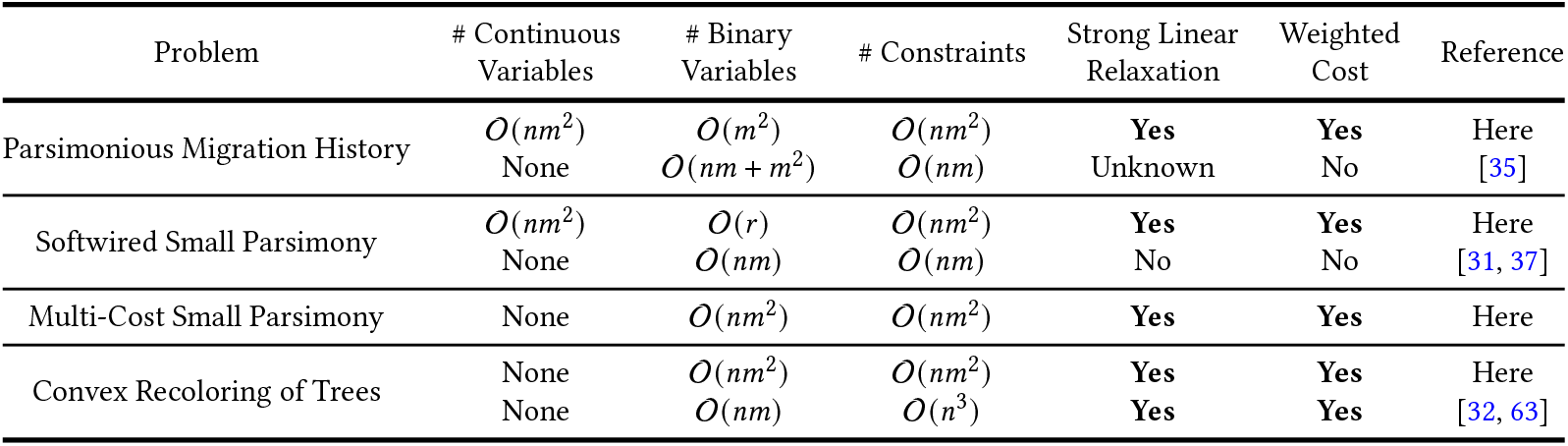
Comparison of the mixed integer programming formulations derived using the tree labeling polytope in this work and existing mixed integer programming formulations. Here, *n* denotes the number of edges in 𝒯 or 𝒩, *m* denotes the size of the label set Ω, and *r* = |*R* (𝒩) | is the number of reticulation vertices in the network 𝒩.

### 4.1 The small parsimony problem

The classical algorithm for the small parsimony problem is based on the *Sankoff-Rousseau recurrence* [3–5]. The algorithm, also known as Sankoff’s algorithm, is an 𝒪(*nm*^2^) bottom-up dynamic programming algorithm based off this recurrence which we briefly describe. Let *A* [*u, i*] denote the minimum cost labeling of the subtree of 𝒯 rooted at vertex *u* such that *u* is labeled by *i*. Then,

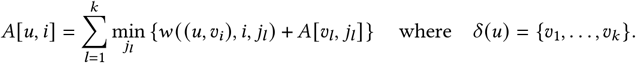

Filling in the table *A*[·, ·], bottom-up leads to the dynamic programming algorithm, where the small parsimony score of the labeling is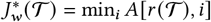. The dynamic programming table *A*[·, ·] has size| *V*(𝒯) |*m* and each entry takes an average of 𝒪(*m*) operations to fill in, leading to a total time complexity of 𝒪(*nm*^2^). When both the tree 𝒯 and cost function *w* are provided as input, this runtime is optimal, as it takes 𝒪(*nm*^2^)*nm*^2^ time to read off the cost function *w*.

The polynomial time algorithm we propose for solving the small parsimony problem is similarly simple. Let 𝒯^′^ be the tree obtained by appending a root *r* ^′^ to 𝒯 and adding the edge (*r* ^′^, *r*). Let *Ax* = *b* be the tree labeling system corresponding to𝒯 ^′^. Using this notation, we obtain the following theorem as a corollary of Theorem 1, which states that the small parsimony problem is solved with a polynomial-sized linear program.

**Corollary 1**. *The small parsimony problem is solved with the linear program* (P) *containing* 𝒪(*nm*^2^) *variables and constraints:*

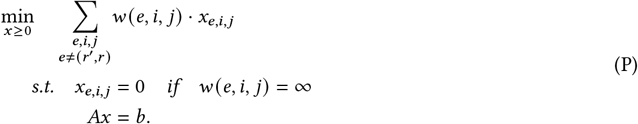

*Proof*. The proof of correctness follows as a corollary from Theorem 1. Specifically, Theorem 1 implies that the linear system *Ax* = *b, x ≥* 0, without the constraint *x*_*e,i,j*_ = 0 if *w* (*e, i, j*) =∞ is totally dual integral. However, the additional constraint consists of replacing the inequality *x*_*e,i,j*_ *≥*0 with the equality *x*_*e,i,j*_ = 0, an operation that preserves total dual integrality (see Proposition 2.3 in [67]). Thus, the above linear program always attains an integral (and in particular, binary) solution (Proposition 1).

We note that in a series of three papers [68–70], Erdős and Székely did provide a *partial* linear programming of the small parsimony problem via a combinatorial duality theorem. However, their formulation did not generalize to rooted trees, or the common case where the cost function is either edge dependent, label dependent, or non-integral. Further, the formulation used 𝒪(*n*^2^*m*^2^) variables and constraints, which is larger by a factor of 𝒪(*n*).

## 5 Results

### 5.1 Fast and accurate migration history inference on simulated data

Using the tree labeling polytope, we derive fastMACHINA, an efficient version of MACHINA, which is an algorithm to infer the most parsimonious history of migrations between a primary tumor and multiple metastases in a cancer patient [11]. fastMACHINA accurately infers migration histories on large simulated instances containing thousands of cells and dozens of anatomical sites (Supplementary Results A.3.1).

We compared fastMACHINA to MACHINA on simulated tumor evolution and metastasis data using a fitness based model from [11] under two different constraints on the migration graph. In particular, we constructed 256 tumor evolution and metastasis scenarios with *n* = 25, 50, 100, 200 extant cells and *e* = 0, 5 lineage tree errors across 16 random seeds, assuming that the migration graph *G* – describing the migrations between different metastatic sites – was either a tree or a directed acyclic graph. The result of each simulated instance was a cellular phylogeny 𝒯, a true migration graph *G*, and a vertex labeling 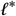 which describes the anatomical location of every cell. Simulation and evaluation details are provided in Supplementary Methods B.1, B.3.

fastMACHINA solved all 256 simulated instances in under 15 minutes, while MACHINA failed to terminate within 24 hours on one of the instances containing *n* = 50 extant cells, half of the instances containing *n* = 100 extant cells, and all of the instances with *n* = 200 or more extant cells (Figure 2a). MACHINA’s runtime was also quite prohibitive, taking a median of over 2 hours to terminate on instances containing only *n* = 100 extant cells (Figure 2a). In contrast,

**Figure 2:**
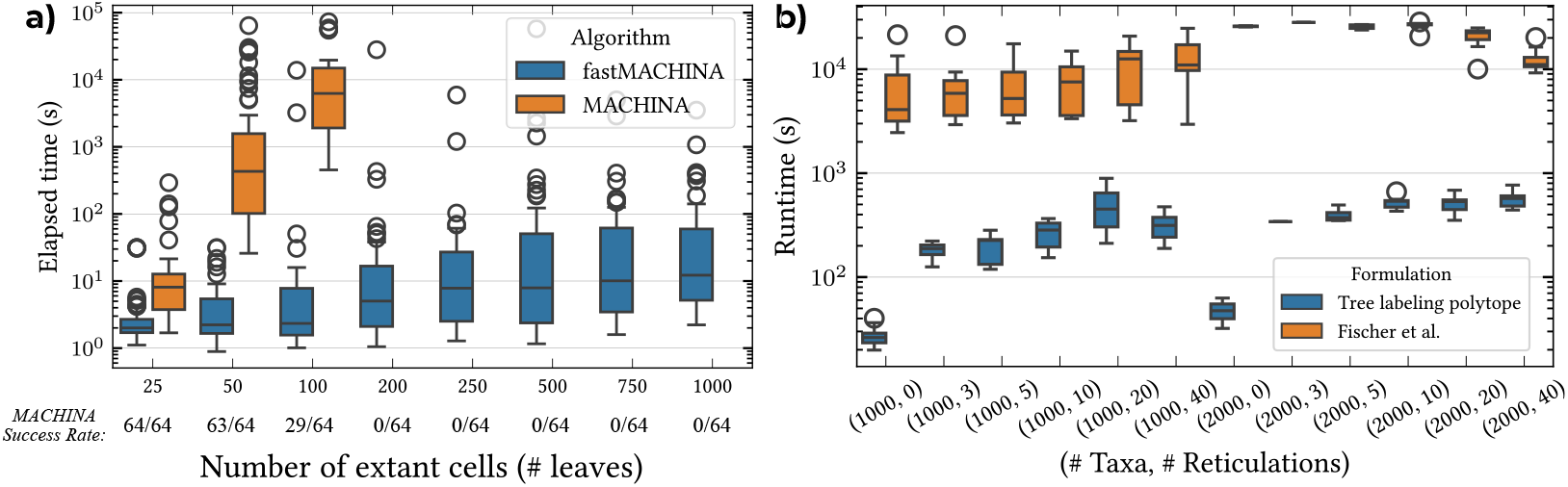
**a)** The runtime of fastMACHINA and MACHINA as a function of the number of extant cells. Each boxplot contains 64 simulated instances and the success frequency of MACHINA within a 24 hour time limit is reported. fastMACHINA terminated on all instances. **b)** The time to solve the mixed integer linear programming formulations of the softwired small parsimony problem versus the number of taxa and reticulation vertices for the tree labeling polytope formulation and the method of Fischer et al. [31].

fastMACHINA easily scaled to instances containing thousands of extant cells, while exhibiting only a slight increase in runtime (Figure 2a). When MACHINA did terminate, fastMACHINA and MACHINA inferred identical migration graphs and parsimony scores.

We compared fastMACHINA to Fitch parsimony, which places no constraints on the inferred migration graph 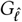, on 256 larger tumor evolution and metastasis scenarios containing *n* = 250, 500, 750, 1000 extant cells where we were unable to apply MACHINA. These tumor evolution and metastasis scenarios contained anywhere from 3 to 71 (median 12) anatomical sites, representing a diversity of scenarios (Supplementary Figure 1). In the absence of any lineage tree error, both fastMACHINA and Fitch parsimony performed well, identifying the ground truth migration graph (Supplementary Figure 4). In the presence of even a small amount of lineage tree error, however, fastMACHINA more accurately recovered the true migration graph in terms of the F1 score between the edges of the true and inferred migration graphs (Supplementary Figure 5). In particular, fastMACHINA found far fewer false positive migration relationships than Fitch parsimony, though found slightly more false negative relationships (Supplementary Figures 2, 3).

### 5.2 Fast softwired small parsimony for simulated phylogenetic networks

We compared our tree labeling polytope based mixed integer programming formulation of the softwired small parsimony problem (Supplementary Results 4.1) on phylogenetic networks to the formulation of Fischer et al. [31], further studied in [37].

In contrast to Fischer et al. formulation, our tree labeling polytope formulation blends the power of mixed integer programming with the tree structure exploited by specialized dynamic programming algorithms [27, 28, 30, 59], a combination that – to our knowledge – is not exploited by any existing algorithm.

We compared our formulation to the Fischer et al. formulation on 198 simulated phylogenetic networks with *n* = 500, 1000, 2000 extant taxa and *r* = 0, 3, 5, 10, 20, 40 reticulations across *s* = 1, …, 11 random seeds using an amino acid alphabet (i.e. |Ω| = 20). To construct each network, we first simulated phylogenetic trees with *n* taxa using the simulator *Ngesh* [71] and uniformly at random constructed *r* reticulation vertices. Then, we uniformly at random sampled the character-state of each extant site from the amino acid alphabet to obtain the sequences at the leaves. To ensure a fair comparison, we benchmarked both our formulation and the Fischer et al. formulation using the mixed-integer programming software Gurobi [72]. Simulation details are further described in Supplementary Methods B.2.

The tree labeling polytope based formulation was substantially faster than the Fischer et al. formulation on all 198/ 198 instances, with the Fisher et al. formulation terminating within an 8 hour time limit on only 171 / 198 instances (Figure 2b). Even in the absence of any reticulation in the network (*r* = 0) – where the problem reduces to small parsimony on trees – the Fischer et al. formulation struggled, taking a mean of 250 times longer to terminate compared to the tree labeling polytope formulation (Figure 2b). Furthermore, while the time required to solve our tree labeling polytope formulation was stable in the size of the network, the time required to solve the Fischer formulation increased considerably in the size of the network (Figure 2b). This is probably, in part, due to the fact that the number of binary variables in our formulation is independent of the size of the network, which is not the case with the Fischer et al. formulation. To quantify the strength of each formulation, we computed the integrality gap 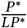 across all instances, where *P*^∗^ is the true objective and *LP* ^∗^ is the value of the linear programming relaxation. While the mean integrality gap of the tree labeling polytope formulation was 1.02, the mean integrality gap for the Fischer et al. formulation was 1.18, demonstrating the improved strength of the formulation obtained using the tree labeling polytope (Supplementary Figure 6).

### 5.3 Large-scale migration history inference on lineage tracing data

We evaluated fastMACHINA on a large single-cell sequencing dataset modeling metastatic dissemination in mice [74]. Using recently developed lineage tracing technology [55, 75, 76], Yang et al. [74] equipped 5,000 human lung adenocarcinoma cells (A549) with a lineage tracer and surgically implanted the cells into the left lung of the immunodeficient mouse “M5k”. Fifty-four days post-surgery, the mouse was sacrificed and tumor samples were collected from the six anatomical sites (Figure 3a), undergoing single-cell sequencing. The result of the experiment was then a set of 41,847 single-cell profiles measured across the six anatomical locations, which were used to reconstruct the cellular lineages of 100 distinct tumor clones. Using the resultant cellular lineages, a key goal of [74] was to infer the routes of metastasis, or in our terminology, the migration graph *G* associated with the lineage tree𝒯. The existing analysis, however, found *G* by selecting a most parsimonious ancestral labeling 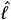 and taking the induced migration graph 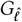. It is known that migration graphs inferred this way can be misleading [11], and that additional constraints are necessary. To resolve this issue, MACHINA [11] imposes additional constraints on the induced migration graph 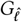 (Section 2). Unfortunately, MACHINA is unable to scale to these large lineage trees, which contain upwards of 5, 616 cells.

**Figure 3:**
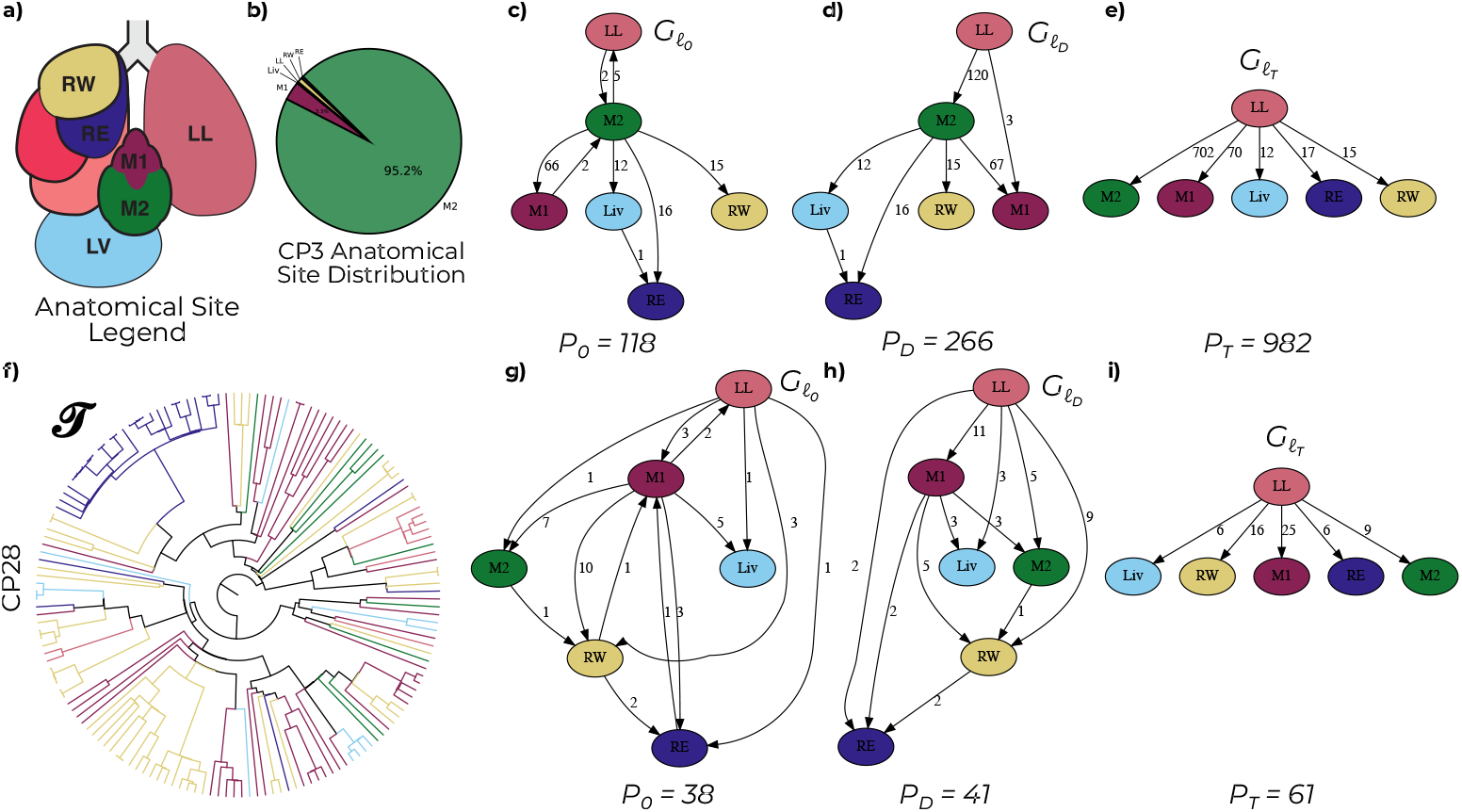
**a)** The 41,847 sequenced cells of mouse “M5k” [15] are located in the left lung (LL), liver (LV), mediastinal lymph tissue (M1, M2), and right lung (RW). **b)** The distribution of the anatomical locations of the 5,616 cells in the largest clone CP3. **c-e)** The migration graphs 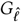 and parsimony scores *P*_0_, *P*_*D*_, *P*_*T*_ induced by the most parsimonious ancestral labeling such that 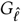 is **c)** unconstrained, **d)** a directed acyclic graph, and *e)* a tree for the lineage tree constructed on clone CP3. **f)** The cell lineage tree constructed by LAML [73] for clone CP28. **g-i)** The migration graphs 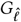 and parsimony scores *P*_0_, *P*_*D*_, *P*_*T*_ induced by the most parsimonious ancestral labeling for clone CP28.

Following [15], we selected the 83 /100 clones from the mouse “M5k” which passed quality control checks and were labeled CP1 through CP100. For each of the 83 clones, we obtained pre-processed character matrices and the anatomical labelings of the cells from the original publication [15]. To infer the cellular lineage trees, we first applied Startle [77], a specialized tool for inferring cellular lineages from CRISPR-Cas9 based lineage tracing data. Next, we inferred branch lengths on these trees and further optimized the topologies using LAML [73].

We applied fastMACHINA (Supplementary Section A.3.1) to infer migration graphs for the 83 lineage trees under three topological constraints on the induced migration graphs. In particular, we inferred the most parsimonious ancestral labelings 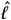 such that the induced migration graph 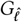 was *i)*unconstrained, *ii)* a directed acyclic graph, and *iii)* a tree. From this, we obtained three ancestral labelings and migration graphs for each of the 83 lineage trees, corresponding to a nested set of constraints on 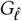. For a fixed tree, we denote the parsimony scores and migration graphs inferred by our version of MACHINA under the three constraints *i), ii), and iii)* as *P*_0_, *P*_*D*_, *P*^*T*^ and *G*_0_, *G*_*D*_, *G*^*T*^ respectively.

For the largest clone CP3, which contained 5,616 cells drawn from 6 anatomical locations, we noticed that the parsimony scores increased substantially (*P*_0_≪ *P*_*D*_≪ *P*^*T*^) as more constraints were added (Figure 3c-e). In particular, an additional 148 metastatic migrations were required to explain the anatomical locations of the cells when no reseeding was allowed to occur. Looking at the inferred migration graphs (Figure 3c-e), we observe that the majority (118/148) of these additional migrations occurred from the left lung to the mediastinal lymph tissue. Since we know the tumor originated in the left lung, a possible explanation is that several cells metastasized to the lymph tissue, upon which a clonal expansion occurred. This would explain both the large number of tumor cells in the lymph tissue (Figure 3b) and the reseeding events back to the left lung. Further, it would explain the increase in the parsimony score (*P*_*D*_ ≪ *P*^*T*^) upon requiring that each site is seeded by a single parent, as it contradicts the hypothesis that both the left lung and the mediastianal tissue act as primary seeding sites.

For the smaller clone CP28, which contained 180 cells drawn from 6 anatomical sites (Figure 3f-i), only an additional three migrations were required to explain the anatomical locations of the cells when reseeding was not allowed to occur (*P*_0_ ≈ *P*_*D*_), not allowing us to reject the hypothesis that reseeding did not occur. Further, one of the reseedings inferred in *G*_0_ occurred from the right lung back to the mediastinal tissue, a surprise given that most of the right lung cells form a single clade in 𝒯 (Figure 3f). Upon enforcing the constraint that each site is seeded by a single parent, however, many more migrations are required (*P*_*D*_ ≪ *P*^*T*^) and all migrations occur from the left lung. Given the reported role of lymph tissue as a transit hub for metastasis [78, 79] and the large increase in migrations, it seems implausible that each parent was seeded by only a single parent in clone CP28.

Performing an analysis across all 83 clones, we analyzed the parsimony scores *P*_0_, *P*_*D*_, *P*^*T*^ for the fifteen largest and the fifteen smallest clones. Immediately, we observed that while the parsimony scores for the largest clones differed across the migration constraints (*P*_0_ ≠ *P*_*D*_ ≠ *P*^*T*^), for the smallest clones the parsimony scores stayed relatively constant (*P*_0_≈ *P*_*D*_≈*P*^*T*^) (Supplementary Figures 7, 8). To quantify the relationship between clone size and parsimony score, we regressed the three parsimony gaps *P*^*T*^− *P*_*D*_, *P*^*T*^ −*P*_0_, and *P*_*D*_− *P*_0_ against the number of cells in the tree (Supplementary Figure 9). Interestingly, we found a strong linear relationship in all three settings (*R*^2^ = 0.99, 0.98, 0.86), suggesting that the size of the tree strongly impacts the parsimony score. Of the many potential explanations, the observed behavior could be due to the increased difficulty in reconstructing larger cellular lineages. If the quality of constructed lineages degrades with its size, it would naturally follow that the anatomical labels are more randomly dispersed across the leaves, increasing the parsimony score directly in relation to the size of the tree. In this experimental setting, this could very much be the case, as the number of CRISPR-Cas9 target sites, which are used as phylogenetic markers, is fixed and independent of the tree size [55].

### 5.4 A quantitative test for the monoclonal theory of metastasis via convex recolorings

We applied our tree labeling polytope based algorithm for the convex recoloring problem (Supplementary Results A.3.4, A.4) to test the monoclonal theory of metastasis [13] on the 83 lineage trees from [55]. The monoclonal theory of metastasis states that each metastasis is founded by a single-cell [13]. Unfortunately, until recently it has been challenging to quantitatively test this hypothesis on real data, as bulk and single-cell DNA sequencing do not provide the necessary phylogenetic resolution.

To construct a quantitative test for the monoclonal theory of metastasis, we answer the following question: *what is the minimum number of cell deletions required to obtain a monoclonal explanation of the cellular lineage?* To formalize this question, we first observe that metastasis is monoclonal if and only if the total number of migrations in the true ancestral labeling *ℓ*^∗^ is *m* − 1, or one less than the number of anatomical sites. Said another way, *ℓ*^∗^ is monoclonal if and only if it is convex on 𝒯. Formally, we then perform this test by determining the minimum number of leaf deletions in 𝒯 such that the leaf labeling on the resulting tree is convex.

We applied the aforementioned test to the 83 lineage trees constructed from [55] by solving the convex recoloring problem on each of the 83 lineage trees (Supplementary Results A.3.4). As a result, for each lineage tree we obtained the minimum number of cell deletions required for a monoclonal explanation along with the deleted cells. Across all 83 lineage trees, a mean of 19.85% of the cells needed to be deleted in order to obtain a monoclonal explanation of the observed lineage (Supplementary Figure 13). Consequently, unless we believe that a large fraction of the cells in the inferred lineage trees are incorrectly placed, there is little support for the monoclonal theory of metastasis across the 83 lineage trees. This conclusion is concordant with an increasingly large body of work on the polyclonality of tumors [56] as well as the notion of *metastatic potential* described in [55].

## 6 Discussion

We introduced the *tree labeling polytope*, a technical tool for parsimonious ancestral labeling problems. As the geometric analogue to Sankoff’s algorithm, the tree labeling polytope enables efficient optimization over ancestral labelings. We anticipate applications of the tree labeling polytope to a larger array of biological questions, such as to estimating the extent of hybridization in bacteria [80, 81] or to inferring cell fate differentiation maps in developing systems [26]. While the tree labeling polytope can be used to find the most probable ancestral labeling by a log-transformation of the probabilities into an edge-dependent cost function, maximum likelihood estimation marginalizes over all ancestral states using Felsenstein’s pruning algorithm [82]. An interesting open question is whether polyhedral techniques can be useful in this setting.

## 7 Acknowledgements

This research is supported by National Cancer Institute (NCI) grants U24CA248453 and U24CA264027 to B.J.R.

## A Supplementary results

We will make use of the following additional definitions and results from polyhedral theory [61]. A *supporting hyperplane* of a polyhedron *P* is a set *H* = {*x* : *c*^*T*^ *x* = *δ*} for some vector *c* and *δ* = max_*x∈ P*_ *c*^*T*^ *x*. A *face* of a polyhedron *P* is the intersection of *P* with a supporting hyperplane. A *facet* of a polyhedron *P* is a maximal face of *P* not equal to *P*. A *vertex* of a polyhedron *P* is a face consisting of a single point. An integer matrix *A* is *totally unimodular* if every submatrix of *A* has determinant in {0, ±1}.

### Proposition 2

([61]). *A vector x is a vertex of P if and only if x is an extreme point of P*.

### Proposition 3

([61]). *Let A be a rational, totally unimodular matrix and b be an integral vector. Then, the polyhedron P* = {*x* : *Ax* = *b, x ≥* 0} *is integral*.

### A.1 Total dual integrality of the tree labeling system

In this section, we provide a proof of Theorem 2 by demonstrating the total dual integrality of the tree labeling system. While for many polytopes, such as flow polytopes, it is simpler to prove integrality by demonstrating the total unimodularity of the corresponding constraint matrix, the tree labeling matrix *A* is not totally unimodular (Supplementary Results A.2). To prove total dual integrality, we develop a polynomial-time dynamic programming algorithm for solving the dual of the linear program min {*c*^*T*^ *x* : *Ax* = *b, x≥* 0}. As a consequence of the analysis of our algorithm, we prove the total dual integrality of the tree labeling system.

To fix notation, let 𝒯 = (*V*(𝒯), *E*(𝒯)) be a rooted tree with root *r* = *r*(𝒯), leaf set *L*(𝒯), and label set Ω = {1, …, *m*}. Let *Ax* = *b, x ≥* 0 be the tree labeling system corresponding to 𝒯. Let min {*c*^*T*^ *x* : *Ax* = *b, x ≥* 0} be the *tree labeling linear program* where the cost vector *c* is assumed to be integral.

To start, we derive an explicit form of the dual problem max{*b*^*T*^*y* : *y*^*T*^ *A* ≤ *c*}, as stated below.

#### Lemma 1.

*The dual linear program* max {*b*^*T*^*y* : *y*^*T*^ *A* ≤ *c*} *is equivalent, up to an additive integer constant, to the following optimization problem:*

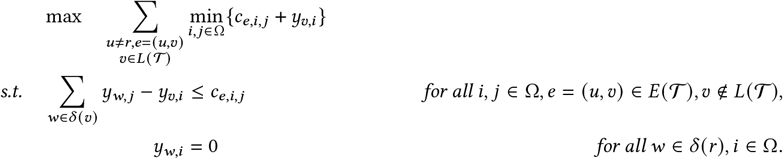

*Further*, max {*b*^*T*^*y* : *y*^*T*^ *A* ≤ *c*} *is attained at an integral vector if the optimum of the above linear program is attained at an integral vector*.

*Proof*. Associate the dual variables *y*_*w,i*_ with the linear constraints (5) and the dual variables *z*_*𝓋*_ with the linear constraints (6). Taking linear combinations of the dual variables with the constraints (5)-(6) yields

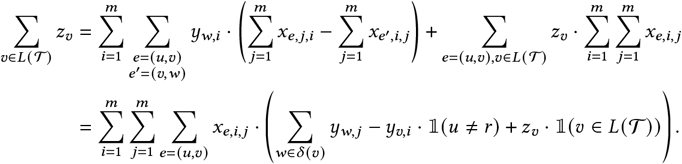

The reasoning for the above equality is that *x* _(*u,𝓋*),*i,j*_ appears positively with *y*_*w,j*_ for all children *w ∈ δ* (*𝓋*), negatively with *y*_*𝓋,i*_, and positively with *z*_*𝓋*_ if *𝓋* is a leaf. Further, if *u* is the root vertex, *x* _(*u,𝓋*),*i,j*_ does not appear negatively with any dual variable. Consequently, the dual linear program becomes:

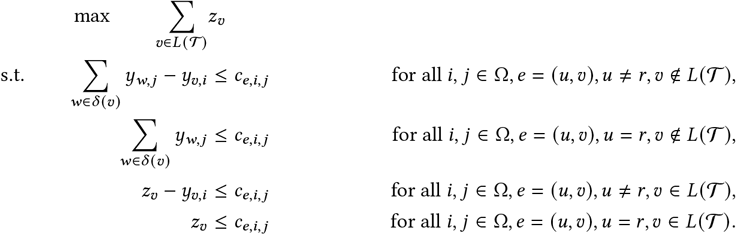

Since we are maximizing with respect to *z*_*𝓋*_ and *z*_*𝓋*_ ≤ *c*_*e,i,j*_ +*y*_*𝓋,i*_ · 𝟙 (*u* ≠ *r*), in every optimal solution *z*_*𝓋*_ = min_*i,j*_ {*c*_*e,i,j*_+ *y*_*𝓋,i*_ · 𝟙 (*u* ≠ *r*)}. If *u* = *r, z*_*𝓋*_ = min_*i,j*_ {*c*_*e,i,j*_} is a fixed integer constant with respect to the optimization objective and can be ignored. Consequently, we optimize out the dual variable *z*_*𝓋*_ to obtain the result. ¦

Next, we state a simple algebraic identity needed in the subsequent analysis.

#### Lemma 2.

*For all real numbers a*_*i,j*_ *and b* _*j*_ *where i ∈* {1, …, *n*} *and j ∈* {1, …, *m*},

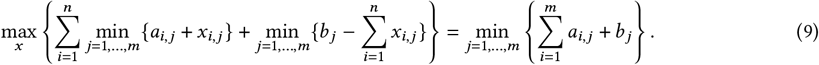

*Proof*. Since the sum of minimums is upper bounded by the minimum of sums, we have that the right-hand side of is an upper bound on the left-hand side of (9). To obtain equality, it then suffices to provide a solution *x* where equality holds. Setting

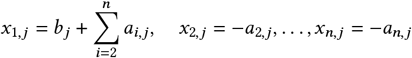

for *j* = 1, …, *m* proves the claim.

We now provide a recursive procedure to compute the optimal value of the dual optimization problem. Then, after backtracking to obtain the dual variables, we observe that the optimal value of the dual linear program max{*b*^*T*^*y* : *y*^*T*^ *A* ≤ *c*} is attained at an integral vector.

#### Lemma 3.

*The optimal value of the dual linear program γ* ^∗^ ≜ max{*b*^*T*^*y* : *y*^*T*^ *A* ≤ *c*} *can be computed in* 𝒪(*nm*^2^) *time and is an integer. There exists an optimal dual vector y*^∗^ = arg max{*b*^*T*^*y* : *y*^*T*^ *A* ≤ *c*} *that is integral and can be computed in* 𝒪(*nm*^2^) *time*.

*Proof*. Let 𝒯_*w*_ be the subtree of 𝒯 rooted at vertex *w*. Consider the following linear program, parameterized by a vertex *l ∈ V* (𝒯) and an *m*-dimensional vector α:

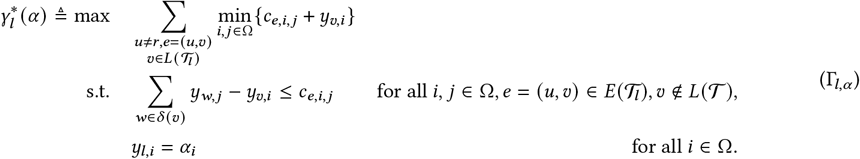

Then, 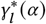 is the optimal value of the dual linear program for the subtree of 𝒯 rooted at vertex *l* such that the dual vector *yl* = α. Consequently, it follows from Lemma 1 that 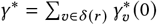 is the value of the optimal solution to the dual linear program. Notice that 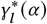 is monotonically increasing in the coordinates α_*i*_ : as α_*i*_ increases, *yl*_,*i*_ increases, which only makes it easier to satisfy the constraints in (Γ*l*,α).

Now, we demonstrate how to compute 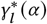 recursively when provided 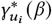 for the children *u*_*i*_ *δ∈* (*l*). From this recurrence, we will obtain a polynomial time dynamic programming algorithm to compute 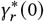, upon which we perform backtracking to recover a set of optimal dual variables.

Let *δ* (*l*) = {*u*_1_, …, *u*_*k*_} be the children of *l* in 𝒯. Then, 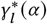 satisfies the following recurrence relation at all non-root vertices *l* with *e* = (*ρ* (*l*), *l*):

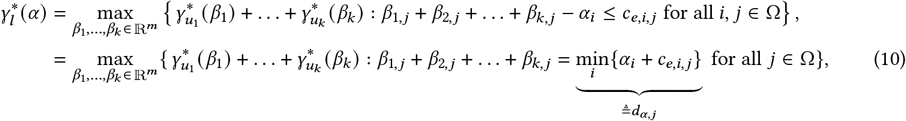

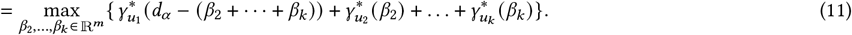

The first equality follows from the definition of (Γ*l*,α). The second from the fact that 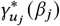is monotonically increasing in the coordinates of β _*j*_. The third follows from the equality constraint in the maximization.

Using this recurrence, we show how to compute 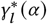 inductively. We say that the inductive hypothesis holds at a vertex *l* if 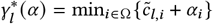 for some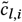.

For the base case, observe that if *l* is a leaf with pendant edge *e*, then 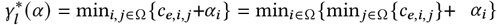. The inductive hypothesis then holds at leaf vertices by setting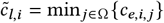.

For the inductive step, let *l* be a non-root vertex with *e* = (*ρ* (*l*), *l*) and assume the inductive hypothesis holds at all descendants of *l*. Plugging the inductive hypothesis into (11) yields,

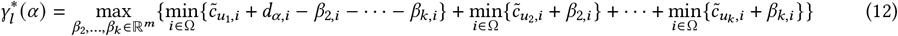

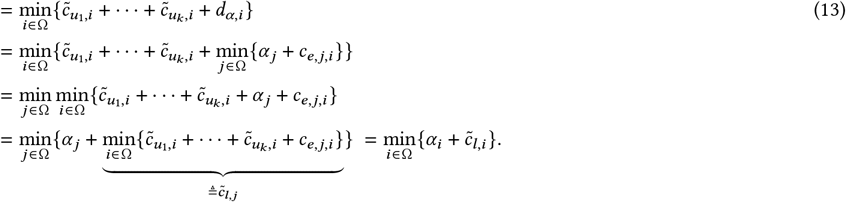

The first equality follows from the induction hypothesis. The second from the algebraic identity provided by Lemma 2. The third from the definition of *d*α_,*j*_. The final equalities follow by algebraic manipulation and setting 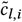 appropriately. Thus, the inductive claim holds at all internal (non-root) vertices.

The above proof that 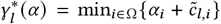also provides a formula for computing 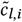 which we turn into a dynamic programming algorithm, described as follows.

1. Visit the vertices in their post-order depth-first search traversal index order (bottom-up).
2. At a leaf vertex *𝓋*, set 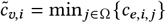 where *e* is the unique edge into *𝓋*.
3. At an internal vertex *𝓋* ≠ *r*, set 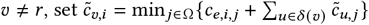where *e* is the unique edge into *𝓋*.
4. Compute the optimal objective value 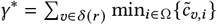.

The above algorithm takes 𝒪(*nm*^2^) time and 𝒪(*nm*) space to compute and store all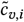, since each value 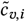 takes an average of 𝒪(*m*) time to compute. From this, we also see that γ ^∗^ takes an integer value, since it is a sum over values *c*_*e,i,j*_, all of which are integral.

The integrality of γ ^∗^, however, is not enough to complete the proof. Indeed, we also need to find an integral set of dual variables *y*_*𝓋,i*_ which realize this objective. Backtracking over the pre-computed values 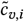 provides a solution, described as follows.

1. Visit the vertices in their pre-order breadth-first search traversal index (top-down).
2. At the root *r*, set *y*_*r,i*_ = 0 for all *i ∈* Ω.
3. At an internal vertex *𝓋*, let *δ* (*𝓋*) = {*u*_1_, …, *u*_*k*_}. Then, set 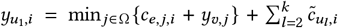, and set 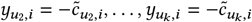 where *e* is the unique edge into *𝓋*.

To see feasibility of the resulting dual variables, note that the constraints are satisfied (and tight) for the dual problem by a straightforward calculation. To see optimality, observe that to obtain (11) from (10), we set *β*_1,*j*_ = *d*α_,*j*_ −(β_2,*j*_ +· · ·+β_*k,j*_) and to obtain (13) from (12) we set 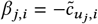 (see proof of Lemma 2). Noting that β_1_, …, β_*k*_ play the role of 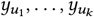and α plays the role of *yl*, this corresponds exactly to the update in our backtracking algorithm. Consequently, the update is an optimal setting of the dual variables 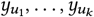, provided we then optimally set the dual variables corresponding to the descendants of *u*_1_, …, *u*_*k*_. Applying this reasoning recursively proves the optimality of our backtracking procedure.

Consequently, after performing backtracking, the dual variables are feasible and achieve the objective γ ^∗^. Since γ ^∗^ is optimal, we have found an integral vector *y* realizing the dual objective. This completes the proof. ¦

As a corollary to the above theorem, we obtain our first main result.

#### Theorem 1.

A few comments are now in order. First, we note the appearance of *self-duality* in our proof of the total dual integrality of the tree labeling system. In particular, interpret 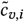 as the minimum cost labeling of the subtree rooted at *w such that the parent of w is labeled by i*. Then, reasoning has that 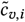satisfies the recurrence relation 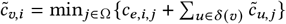. Interestingly, however, this is exactly how we compute the dual objective γ ^∗^ in the preceding proof. Thus, the dual is essentially solved by a modified version of Sankoff’s algorithm, leading us to refer to the tree labeling system as self-dual.

#### Theorem 2.

*Given a rooted tree*𝒯 *and a label set* Ω, *let Ax* = *b, x≥* 0 *be the tree labeling system corresponding to*𝒯 ^′^. *Then, the tree labeling polytope is*

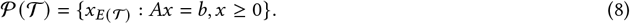

*Proof*. To complete the proof, we need to show two inclusions hold. The first inclusion that

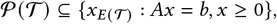

is straightforward. Consider a vertex labeling 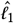 of 𝒯 and an arbitrary extension 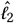of 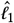 to a vertex labeling of 𝒯 ′. Since 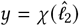 satisfies all inequalities (5)-(7), it follows that *Ay* = *b, y* ≥ 0. Since by definition 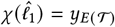, we know that 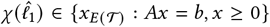. Since this holds for an arbitrary vertex labeling, it holds for all vertex labelings, completing this direction of the proof.

For the second inclusion, let *y* be a vertex of the polytope {*x*_*E*(𝒯)_ : *Ax* = *b, x≥* 0}. By the total dual integrality of the tree labeling system (Theorem 1) it follows that *y* is an integral vector. Indeed, *y* is also a binary vector. To see this, note that for all pendant edges *e*, Σ _*i,j*_ *y*_*e,i,j*_ = 1, implying *y*_*e,i,j*_ is binary. Propagating this constraint up the tree (using (5)-(6)), however, we also have Σ_*i,j*_ *y*_*e,i,j*_ = 1 for an arbitrary edge *e*, allowing us to also conclude that *y* is binary. Since *y* is a binary vector satisfying (5), each vertex other than *r* ^′^, which is not in 𝒯, has a unique label. Thus, there exists a unique vertex labeling 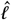 of 𝒯 with 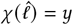), completing the proof. ¦

### A.2 Total unimodularity of the tree labeling matrix

A stronger sufficient condition for proving Theorem 2 would be to to show that the tree labeling matrix *A* is totally unimodular, from which it follows by Proposition 3. Typically, this is simpler than proving total dual integrality, in fact, there is even a polynomial time algorithm for determining the total unimodularity of a matrix. Unsurprisingly, this is not the case, and we provide a simple counter-example to the total unimodularity of the tree labeling matrix. Consider the rooted tree 𝒯 = (*V, E*) with vertex and edge set

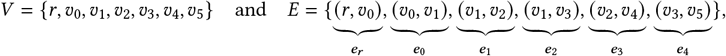

where the root vertex is *r*. The tree labeling matrix *A* associated with 𝒯 also depends on the alphabet Ω, which we set to Ω = {0, 1, 2}. Then, consider the submatrix *A*^′^ of *A* formed by the subset of variables with indices

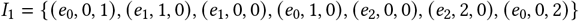

and the subset of constraints from (5) with indices

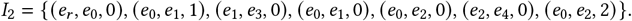

In particular,

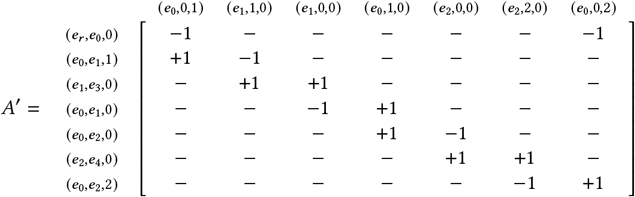

where the columns correspond to the variables indexed by *I*_1_ and the rows the in (5) constraints indexed by *I*_2_. It follows from a computation that det *A*^′^ = −2, a violation of the total unimodularity of *A*. Interestingly, our numerical experiments for the special case when the number of labels *m* = 2 and the 𝒯 tree is binary suggest that that the matrix *A* is totally unimodular in this special case. We state this as a conjecture formally below.

#### Conjecture 1.

*Let* 𝒯 *be a rooted, binary tree and let* |Ω| = 2. *Then, the tree* 𝒯 *labeling matrix A corresponding to is totally unimodular*.

### A.3 Algorithms

#### A.3.1 The parsimonious migration history problem

In the parsimonious migration history problem we possess the additional constraint that the induced migration graph 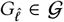. For simplicity of presentation, we will work with in the case where 𝒢 consists of the set of rooted trees, though our techniques also apply to the case where 𝒢 is the set of directed acyclic graphs or directed paths with only slight differences, described briefly at the end of the section. We also assume the inferred vertex labeling 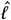 is required to be an extension of the leaf labeling *ℓ* (Section 2.1), as this implies the weak connectivity of 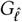, stated as follows.

##### Lemma 4.

*If* 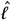 *extends ℓ and* range(*ℓ*) = Ω, *then the migration graph* 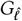 *is weakly connected*.

*Proof*. Let 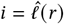 be the label of the root vertex *r* = *r*(𝒯). Consider any other label *j* ≠ *i* ∈ Ω. Since range *ℓ* = Ω, there exists *𝓋 ∈ V*(𝒯) such that *ℓ 𝓋* = *j*. Consider the path *P* = *r, u*_1_, …, *u, 𝓋* in from *r* to *𝓋*. The path *P* yields a walk 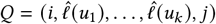, *j* in 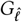. Eliminating tours in *Q* yields a path from *i* to *j* in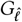. Since there is a path from *i* to any other vertex *j* in 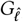, the migration graph 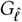 is weakly connected, completing the proof. ¦

To start, we model the edges of the migration graph with *m* (*m*− 1) binary variables *y*_*i,j*_*∈* {0, 1} which denote the presence or absence of an edge *i, j*. To relate the migration graph to the labeling, we add a set of linking constraints *y*_*i,j*_ *≥ x*_*e,i,j*_ for all *e ∈ E*(𝒯), which connect the migration graph to the labeling 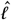 associated with *x*. Specifically, if there is a label change from *i* to *j* across edge *e* then *x*_*e,i,j*_ = 1, which implies that *y*_*i,j*_ = 1. To ensure the migration graph defined by *y*_*i,j*_ forms a tree, there are many options, such as using flow, power set, or spanning tree constraints. Because of the weak connectivity of 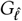 (Lemma 4), however, we enforce that *y*_*i,j*_ is a tree by simply setting Σ_*i,j*_ *y*_*i,j*_ = *m* − 1. In this setting, it is easy to verify that the following integer linear program then solves the parsimonious migration history problem:

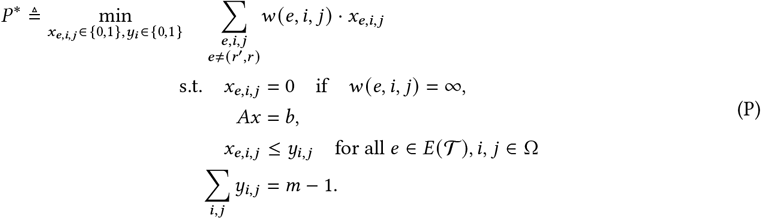

Interestingly, the integrality constraint *x*_*e,i,j*_ *∈* {0, 1} in (P) need not be explicitly enforced and can be relaxed to *x*_*e,i,j*_*≥* 0. This is because for any fixed solution *y*_*i,j*_, the only constraint relating *y*_*i,j*_ and *x*_*e,i,j*_ is *x*_*e,i,j*_ ≤ *y*_*i,j*_. If *y*_*i,j*_ = 0, then this turns into the equality constraint *x*_*e,i,j*_ = 0, which is well-known to preserve total dual integrality (see Proposition 2.3 in [67]). Similarly, if *y*_*i,j*_ = 1, then since *x*_*e,i,j*_ ≤ 1 the constraints are *x*_*e,i,j*_ ≤*y*_*i,j*_ is redundant. Consequently, for any fixed binary vector *y*, the above linear system is totally dual integral, implying that the optimum is always achieved by an integral *x*. Thus, we can replace *P* ^∗^ with the following (equivalent) mixed integer linear program:

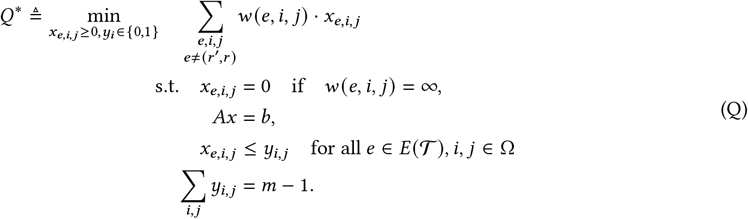

Of note, the number of integer variables in (Q) is of order 𝒪(*m*^2^) which is independent of the size of𝒯. Consequently, we refer to the formulation (Q) as *fixed parameter tractable* in *m*, since it can be solved in polynomial time by solving a linear program for each setting of *y*_*i,j*_. In this way, the formulation (Q) blends together the power of integer programming and Sankoff’s algorithm, resolving the trade-off between integer programming and fixed parameter tractability described in [35].

However, as mentioned earlier, the above formulation (Q) has another nice property related to the strength of its linear programming relaxation, described as follows. We form the Lagrangian relaxation of the above integer program by “convexifying” the complicating constraint. In detail, we form the Lagrangian dual problem by relaxing the constraints associated with *y*_*i,j*_ :

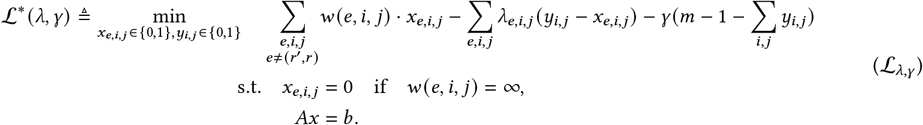

Then, since (ℒλ,γ) is totally dual integral when restricted to *x*, totally unimodular when restricted to *y* (as *y*_*i,j*_*∈* [0, 1] is totally unimodular) and there are no constraints connecting *x* and *y*, the linear relaxation of (ℒλ,γ) always attains its optimum at an integer vector (*x, y*). Using the well-known fact that if there is always an integral solution to the linear relaxation of the Lagrangian relaxation, the value obtained by the linear programming relaxation matches that of the Lagrangian relaxation, we obtain the following result.

##### Corollary 2.

*The parsimonious migration history problem can be expressed as the mixed integer linear program* (Q) *with* 𝒪(*nm*^2^) *continuous variables and* 𝒪(*m*^2^) *binary variables whose linear relaxation*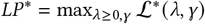.

To extend the above formulation to the case where 𝒢 is the set of dLirected acyclic graphs, one could, instead of enforcing that Σ _*i,j*_ *y*_*i,j*_ = 1, append the cycle elimination constraints Σ_(*i,j*) *∈*_C *y*_*i,j*_ ≤ |C | − 1 for all directed cycles C. This adds an exponential number of constraints, but these can be handled implicitly with a cutting plane approach as there is a polynomial time separation oracle for the cycle elimination constraints [83]. Other, polynomial size formulations for ensuring that *y*_*i,j*_ induces a directed acyclic graph also exist, though we expect the exponential sized formulation together with a separation oracle to be preferable. The case when 𝒢 is a path is a bit easier: a directed graph *G* is a path if and only if it is a tree where no vertex has out-degree greater than or equal to 2. Thus, appending the *m* constraints _*i*_ *y*_*i,j*_ ≤ 2 to (Q) ensures that *y*_*i,j*_ induces a path.

#### A.3.2 The softwired small parsimony problem on phylogenetic networks

In the softwired small parsimony problem, we no longer have a rooted tree 𝒯, but rather a phylogenetic network 𝒩. As before, we append a synthetic root *r* ^′^ to 𝒩 and add the edge (*r* ^′^, *r*) to 𝒩. Noting that the softwired small parsimony score is defined as the minimum over all trees *displayed* by 𝒩, we need additional variables to select the displayed tree. We thus create a binary variable *y*_*e*_ to take value 1 if and only if *e* is in the tree displayed by 𝒩. For all *e* ∉ *E* (ℛ (𝒩)), we fix *y*_*e*_ = 1, ensuring that these edges are always included in the displayed tree. Then, the following integer linear program solves the softwired small parsimony problem on phylogenetic networks:

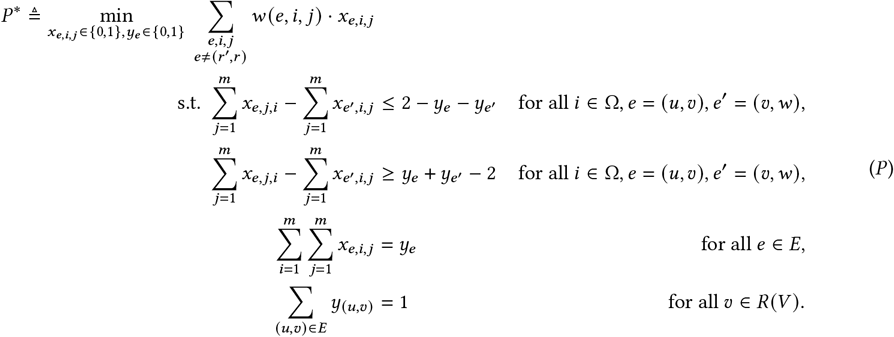

We note that, as opposed to the parsimonious migration history problem (Supplementary Results A.3.1), it is less obvious that (*P*) solves the small parsimony problem on phylogenetic networks. To see that (*P*) is correct, first note that the last constraint in (*P*) implies that the selected edges *F* = {*f ∈E*(𝒩) : *y*_*f*_ = 1} form a rooted spanning tree of 𝒩. For any fixed set of selected edges *F*, the variables *x*_*f, i,j*_ = 0 if *f* ∉ *F*. Further, the first two constraints of (*P*) are redundant if either *e, e*^′^ ∉ *F* and reduce to the matching constraints (5) of the tree labeling system otherwise.

Analyzing the above integer linear program more carefully, we observe that integrality on *x*_*e,i,j*_ need not be explicitly enforced. Thus, we can replace (*P*) with the following (equivalent) mixed integer linear program.

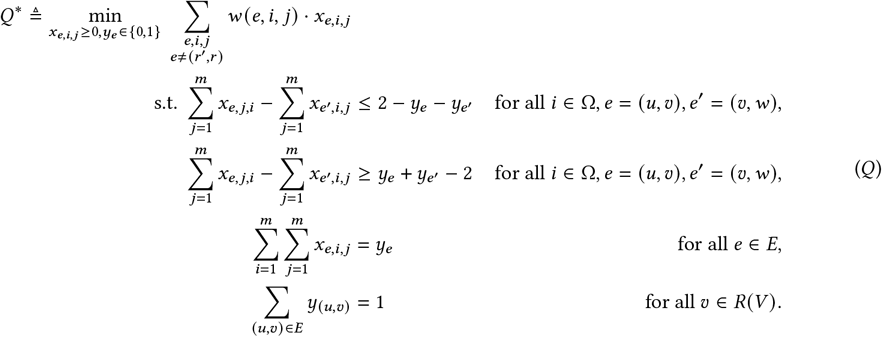

To see that this formulation is correct, we consider any fixing of the binary vector *y* with *F* _′_= {*f ∈ E* (𝒩) : *y*_*f*_ = 1}. By the same reasoning as above, the first two constraints of (*P*) are redundant if either *e, e* ∉ *F* and reduce to the matching constraints (5) of the tree labeling system otherwise. Further, *x*_*f, i,j*_ is set to 0 for all *f* ∉ *F*, not affecting the objective. Since for a fixed binary vector *y* the above reduces to the tree labeling system with additional equality constraints, the resultant system is totally dual integral (Proposition 2.3 in [67]). Consequently, it attains its solution at an integral vector, proving that integrality need not be explicitly enforced on the variables *x*_*e,i,j*_ in (*Q*), leading to the following theorem.

##### Corollary 3.

*The softwired small parsimony problem can be expressed as the mixed integer linear program* (*Q*) *containing* 𝒪(*r*) *binary variables and* 𝒪(*nm*^2^) *continuous variables whose linear relaxation LP* ^∗^ → *Q*^∗^ *as r* → 0.

#### A.3.3 The multi-cost small parsimony problem

Suppose we have two cost functions *w*_1_, *w*_2_ as well as a bound on the second cost *k*. The multi-cost small parsimony problem is nearly a small parsimony problem with cost function *w*_1_, but possesses the additional linear constraint that Σ_*e,i,j*_ *w*_2_(*e, i, j*) · *x*_*e,i,j*_ ≤ *k*. Because of this similarity, it is tempting to simply add this linear constraint to the linear program which solves the small parsimony problem (Section 4.1), obtaining:

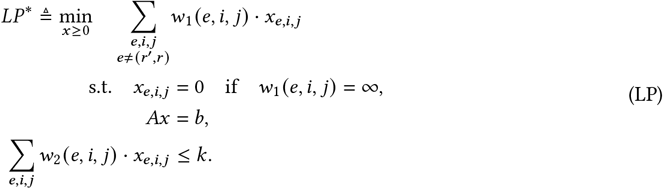

Unfortunately, this additional constraint destroys the integrality of the linear program.

Instead, we suggest that one solves the following integer linear program, obtained by enforcing integrality on *x* :

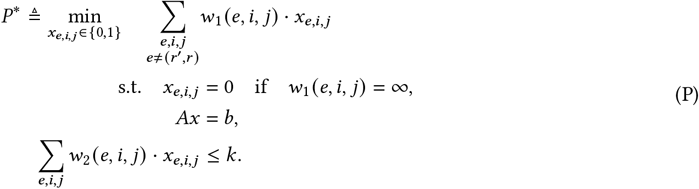

Of course, integer programs are in general difficult to solve – it seems we are back where we started. However, as mentioned earlier, the above formulation (P) has a particularly nice property related to the strength of its linear programming relaxation, described as follows. We form the Lagrangian relaxation of the above integer program by “convexifying” the complicating constraint:

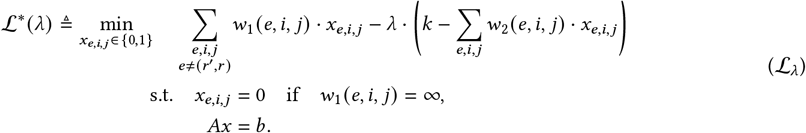

Then, for any *λ >* 0, ℒ^−^ (*λ*) is clearly a lower bound on *P* ^−^. In fact, the relaxation (ℒ_*λ*_) is the optimal solution to the small parsimony problem with weight function 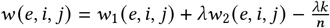. Said another way, we have that 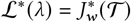 where *w* is defined as the weighted sum of *w*_1_ and *w*_2_.

Thus, we can view the Lagrangian relaxation (ℒ_*λ*_) as either *i*) replacing the hard constraint that ∑_*e,i,j*_ *w*_2_(*e, i, j*) ≤ *k* with the soft linear penalty *λ* · (*k* − ∑_*e,i,j*_ *w*_2_(*e, i, j*) · *x*_*e,i,j*_) or *ii)* solving a perturbed small parsimony problem. The Lagrangian dual problem, then consists of finding the relaxation that maximizes the lower bound ℒ^−^ (*λ*), namely solving the one-dimensional convex optimization problem

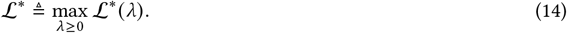

Intuitively, one expects that ℒ^−^ provides a quite reasonable lower bound *P* ^−^, experiments will show this is indeed the case.

It is well known (see Corollary 10.1 [46]) that if there is always an integral solution to the linear relaxation of the Lagrangian relaxation, one obtains ℒ^−^ = *LP* ^−^. Due to the tree labeling polytope, we know that there is always an integral solution to the Lagrangian relaxation, since the linear system in (ℒ_*λ*_) is totally dual integral. Consequently, it follows that ℒ^−^ = *LP* ^−^, or in other words, that the linear programming relaxation of *P* ^−^ is as strong as the Lagrangian relaxation, though in general we only have ℒ^−^ ≥ *LP* ^−^. This is summarized as the following corollary of Theorem 1.

##### Corollary 4.

*The multi-cost small parsimony problem can be expressed as the integer linear program* (P) *with* 𝒪 (*nm*^2^) *binary variables and constraints whose linear relaxation LP* ^−^ = ℒ^−^.

#### A.3.4 The convex recoloring problem on trees

Originally motivated by applications in phylogenetics, the *convex recoloring problem on trees* [84] aims to find the minimum number of leaf label changes such that the resultant labeling is *convex*^5^. The convex recoloring problem is known to be NP-complete on paths, trees, and general graphs [29, 84, 85], and has a long history in discrete mathematics, theoretical computer science, and operations research. Due to its computational intractability, a wide variety of approaches for solving the problem have been applied, spanning parameterized exact algorithms [cite], approximation algorithms [54, 86, 87], integer programming algorithms [32, 33, 63, 88, 89], and heuristics [90].

However, the convex recoloring problem on trees has another interpretation which has yet to be exploited. First, observe that a leaf labeling *ℓ* is *convex* if and only there exists an extension 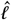 of *ℓ* such that the small parsimony score 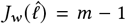 under Wagner parsimony *w e, i, j* = 𝟙(*i* ≠ *j*). Then, it follows that the convex recoloring problem finds the minimum number of leaf label changes such that the small parsimony score is *m* − 1. Following this reasoning further, define *w*_1_ (*e, i, j*) = 1 if *e* = (*u, ν*) is a pendant edge with *j* ≠ *ℓ* (*ν*) and set *w*_1_ (*e, i, j*) = 0 otherwise. Let *w*_2_ (*e, i, j*) = 𝟙(*i* ≠ *j*). Then, the convex recoloring problem is equivalent to the multi-cost small parsimony problem with cost functions *w*_1_, *w*_2_ and *k* = *m* − 1, as *w*_1_ records the number of leaf label changes and *w*_2_ records the Wagner parsimony score.

### A.4 Fast convex recoloring of trees

An advantage of viewing the convex recoloring problem from the perspective of small parsimony is that it enables us to apply the polyhedral tools we developed for the multi-cost small parsimony problem (Supplementary Results A.3.3). To test our application of multi-cost small parsimony to the convex recoloring problem, we simulated 80 leaf labeled phylogenetic trees and applied the integer linear programming formulation developed for the multi-cost small parsimony problem (Section A.3.3). For comparison, we implemented and benchmarked against the mixed integer programmming formulation for the convex recoloring problem described in [32], which is known to be state-of-the-art. Then, we simulated 80 random instances across 4 different conditions using our tumor evolution and metastasis simulator with *e* = 5 errors and migration rate 10^−4^, containing *n* = 50, 100, 150, 200 leaves (Supplementary Methods B.1). For each simulated instance, we measured the strength of the linear programming relaxation, the solver’s runtime, and the minimum number of label changes required to solve the problem.

On all 80/80 simulated instances, the tree labeling polytope based mixed integer linear programming formulation outperformed the state-of-the-art formulation, terminating in under a minute on all instances (Supplementary Figure 10). In contrast, the Campelo formulation [32] only terminated on 68/80 of the instances within a 2 hour time limit, and took around 1000 seconds to terminate on instances containing *n* = 200 cells. While our formulation had slightly weaker linear relaxations than the Campelo formulation, the relaxation objective was always within a constant multiplicative factor of 3 from the true objective (Supplementary Figure 12). Further, the memory usage of the tree labeling polytope formulation was small (*<* 1 GB), whereas the Campelo formulation used up to 32 GB of memory on several instances (Supplementary Figure 11).

## B. Supplementary methods

### B.1 Simulating tumor evolution and metastasis

We simulated tumor evolution and metastasis using the fitness-based model described in [35] which is an extension of the agent model introduced in [91]. The model describes the evolution of cells over a finite number of generations, where associated with each cell is a heritable fitness and a heritable anatomical location. To construct our simulations, we implemented the cell tree simulation procedure described in Supplementary Results A.1 of [35]. Regarding the model parameters, we fixed the driver fitness effect to 10^−1^, the passenger fitness effect to 0, the driver mutation probability to 2 × 10^−7^, the carrying capacity to 5,000, the mutation rate to 10^−1^, and the migration rate to 10^−3^. Additionally, upon a migration event we migrated to a new, as opposed to an existing, anatomical site with probability 1/3 as opposed to the 1/2 used in [35].

With the parameters of the fitness-based model fixed as described above, each simulation was defined by four remaining parameters: the number of extant cells *n*, the number of lineage tree errors *e*, the migration graph class 𝒢, and the random seed *r*. To construct each instance, we simulated tumor evolution using the fitness-based model for 40 generation using the graph class 𝒢 and random seed *r* to obtain a tree 𝒯 labeled by anatomical sites *ℓ* : *V* (𝒯) → Ω. We then took this tree, uniformly sampled *n* leaves from, and took the tree induced on this subset of leaves to obtain 𝒯^′^. Next, we sequentially performed *e* random subtree-prune-and-regraft operations (selected uniformly at random) to obtain a tree 𝒯^′′^. The restriction of *ℓ* to the subset of vertices *V* (𝒯^′′^) ⊆ *V* (𝒯) provided our vertex labeling *ℓ*^−^ and migration graph 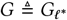. Then our simulated instance was the cellular phylogeny 𝒯^′′^, the migration graph *G*, and the vertex labeling *ℓ*^−^.

### B.2 Simulating phylogenetic networks

Each simulation was described by the number of extant taxa *n*, the number of reticulations *r*, and the random seed *s*. To construct each phylogentic network, we first simulated phylogenetic tree 𝒯 with *n* taxa using the simulator *Ngesh version 1*.*0* [71] with default parameters. Then, we took the partial order ⪯_𝒯_ induced by 𝒯. To construct the network 𝒩, we added *r* reticulation nodes sequentially, resulting in a sequence of networks = 𝒩_0_, 𝒩_1_, …, 𝒩 _*r*_ = 𝒩. To construct 𝒩_*i*_ from 𝒩_*i* − 1_, we uniformly at random sampled a pair of nodes *u* and *ν* in *V* (𝒩_*i* − 1_) such that *u* ≠ *ν*, (*u, ν*) ∉ *E* (𝒩_*i*− 1_), *ν* ∉ *R* (𝒩_*i*− 1_), and *u* ∉ *L* (𝒩_*i*− 1_). Then, if 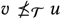 we added the edge (*u, ν*) to 𝒩_*i*− 1_ to obtain 𝒩_*i*_ and otherwise repeated the process until we found such a pair. Since all directed edges added during this process respected the tree order ⪯_𝒯_ it is guaranteed that 𝒩 is acyclic, and thus a phylogenetic network with *r* reticulation vertices of in-degree 2. To construct a leaf labeling *ℓ* : *L*(𝒩) → Ω, we followed the procedure in [31], which uniformly at random assigns leaf labels from Ω = {1, …, 20} to each leaf *ν* ∈ *L*(𝒩). In sum, our simulation procedure resulted in a phylogenetic network 𝒩 and a leaf labeling *ℓ*.

### B.3 Evaluating algorithms for the parsimonious migration history problem

To evaluate MACHINA, we first built and linked MACHINA against version 10.0.3 of Gurobi [72] following the instructions on the GitHub repository github.com/raphael-group/machina. To run MACHINA, we applied the `pmh` mode of the software, which exactly solves the parsimonious migration history problem using mixed-integer linear programming. Using the `pmh` mode of the software, we passed in the cell lineage tree 𝒯, the leaf labeling *ℓ*, and the root label. To evaluate fastMACHINA, we accessed Gurobi version 10.0.3 through the Python API gurobipy. To run fastMACHINA, we passed in the cell lineage tree 𝒯 and the leaf labeling *ℓ* to the `tlp.py` script available on the GitHub repository github.com/raphael-group/tree-labeling-polytope.

For both MACHINA and fastMACHINA, we used the Linux command line utility `/usr/bin/time` to measure the runtime of the software. Both methods were provided with a dedicated 16 core processor, 16 GB of memory, and a 24 hour time limit.

## List of Supplementary Figures

**Supplementary Figure 1:**
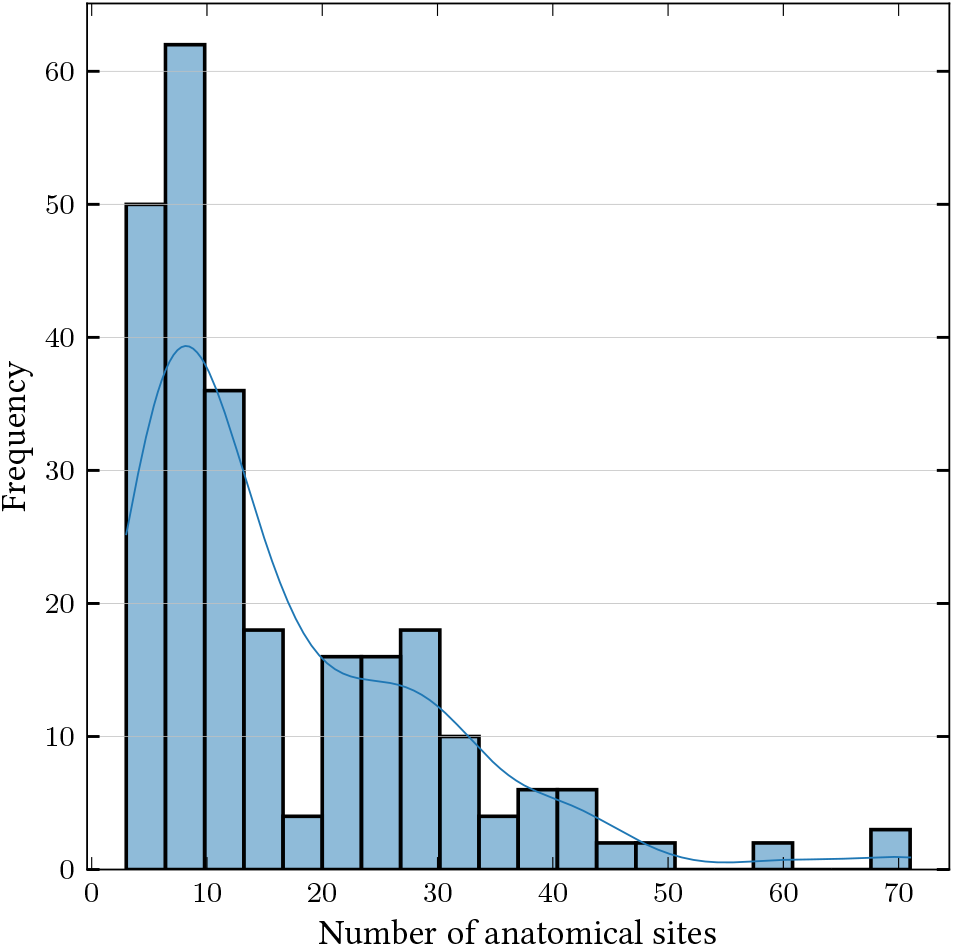
Distribution of anatomical sites in 256 large simulated tumor evolution and metastasis scenarios containing *n* = 250, 500, 750, 1000 extant cells.

**Supplementary Figure 2:**
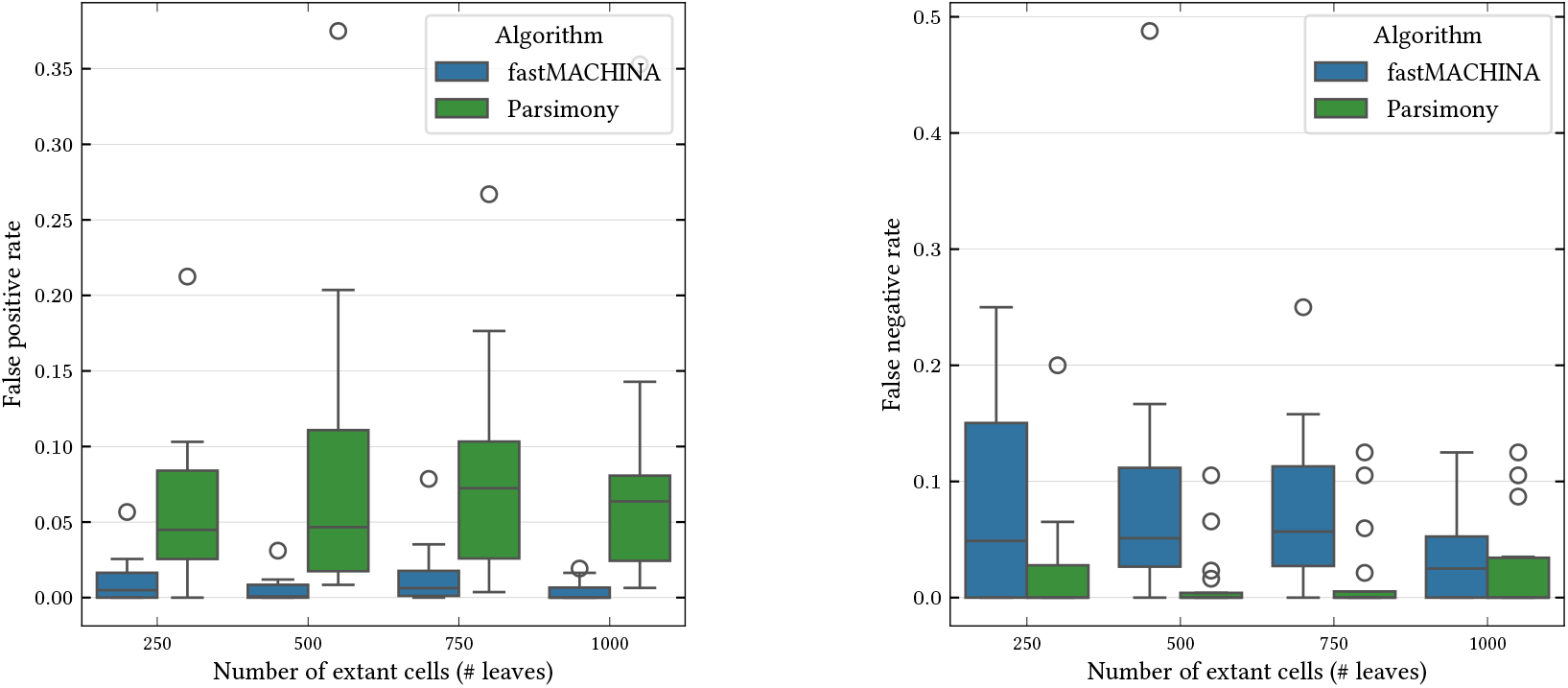
False positive and false negative rates for inferring the edge sets *E*(*G*) of the ground truth migration graph *G* in the presence of *e* = 5 lineage tree errors for the 128 large simulated tumor evolution and metastasis scenarios containing *n* = 250, 500, 750, 1000 extant cells when the ground truth migration graph *G* is a tree.

**Supplementary Figure 3:**
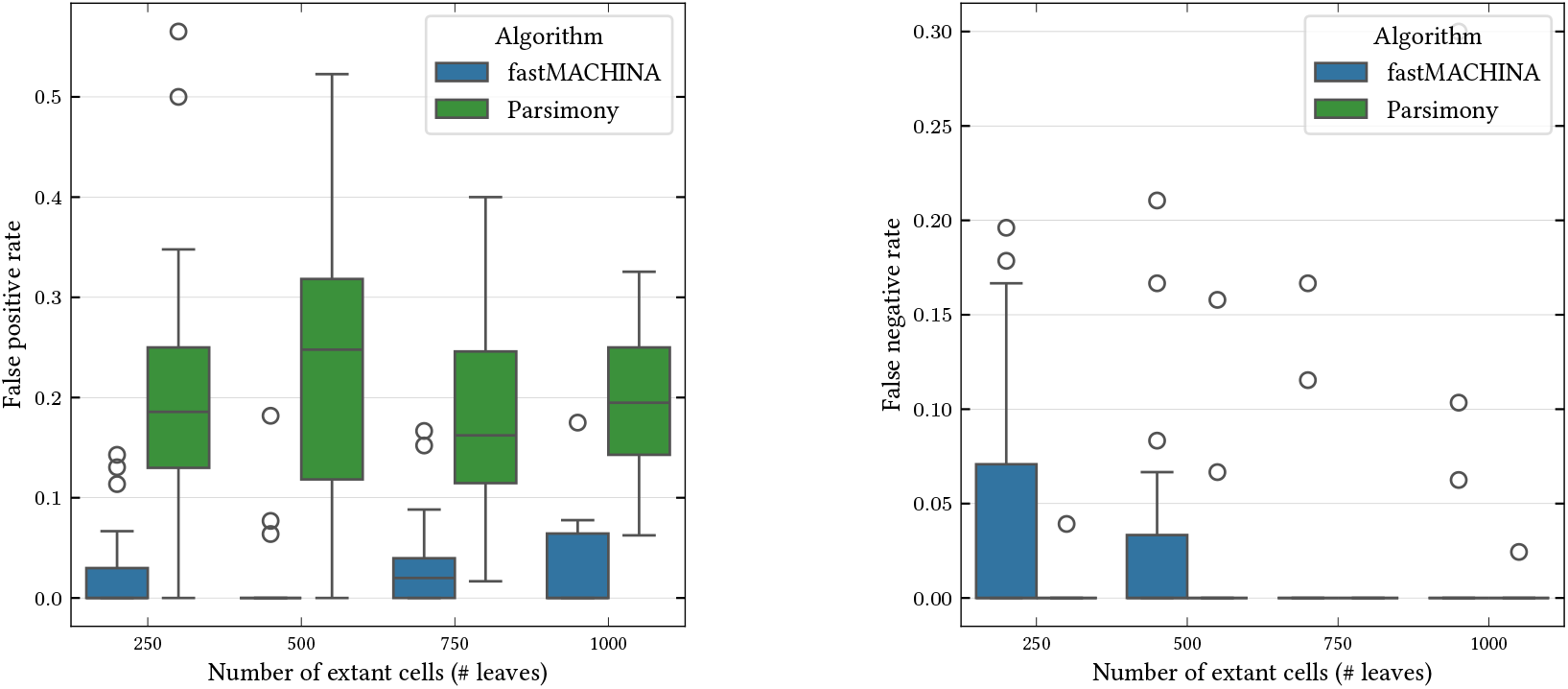
False positive and false negative rates for inferring the edge sets *E* (*G*) of the ground truth migration graph *G* in the presence (*e* = 5) of lineage tree errors for the 128 large simulated tumor evolution and metastasis scenarios containing *n* = 250, 500, 750, 1000 extant cells when the ground truth migration graph *G* is a directed acyclic graph.

**Supplementary Figure 4:**
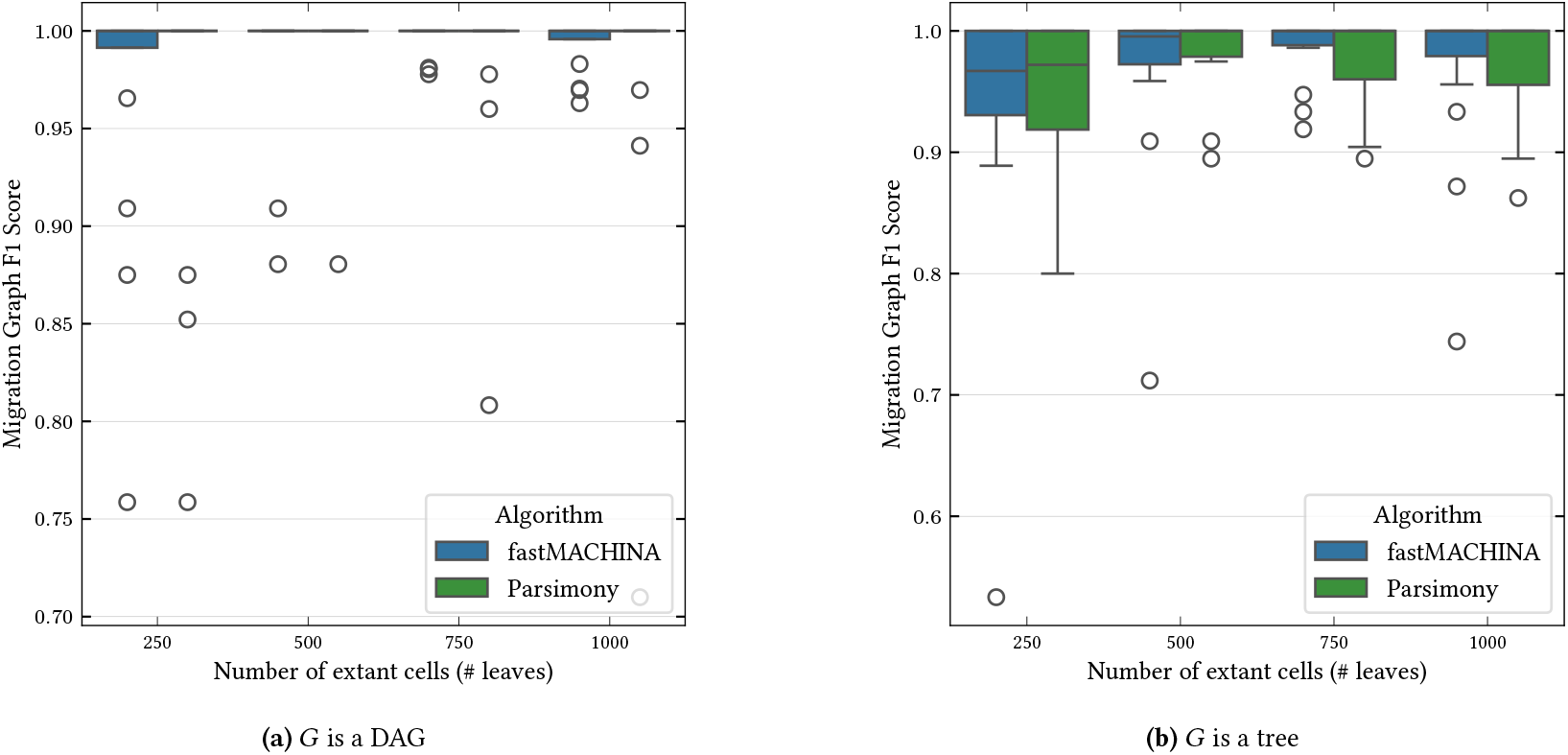
F1 score for inferring the edge sets *E* (*G*) of the ground truth migration graph *G* in the absence (*e* = 0) of lineage tree errors for the 128 large simulated tumor evolution and metastasis scenarios containing *n* = 250, 500, 750, 1000 extant cells when the ground truth migration graph.

**Supplementary Figure 5:**
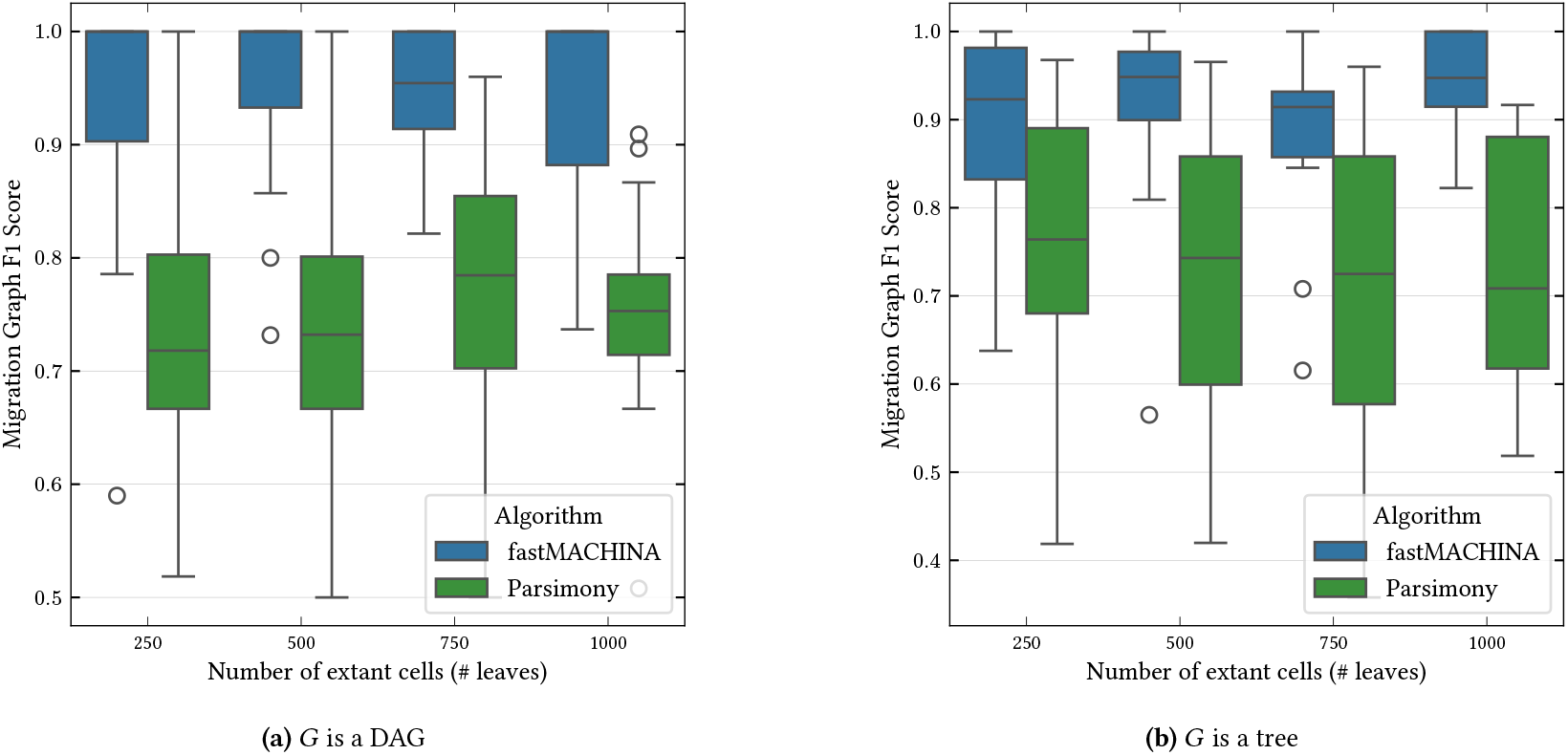
F1 score for inferring the edge sets *E* (*G*) of the ground truth migration graph *G* in the presence (*e* = 5) of lineage tree errors for the 128 large simulated tumor evolution and metastasis scenarios containing *n* = 250, 500, 750, 1000 extant cells when the ground truth migration graph.

**Supplementary Figure 6:**
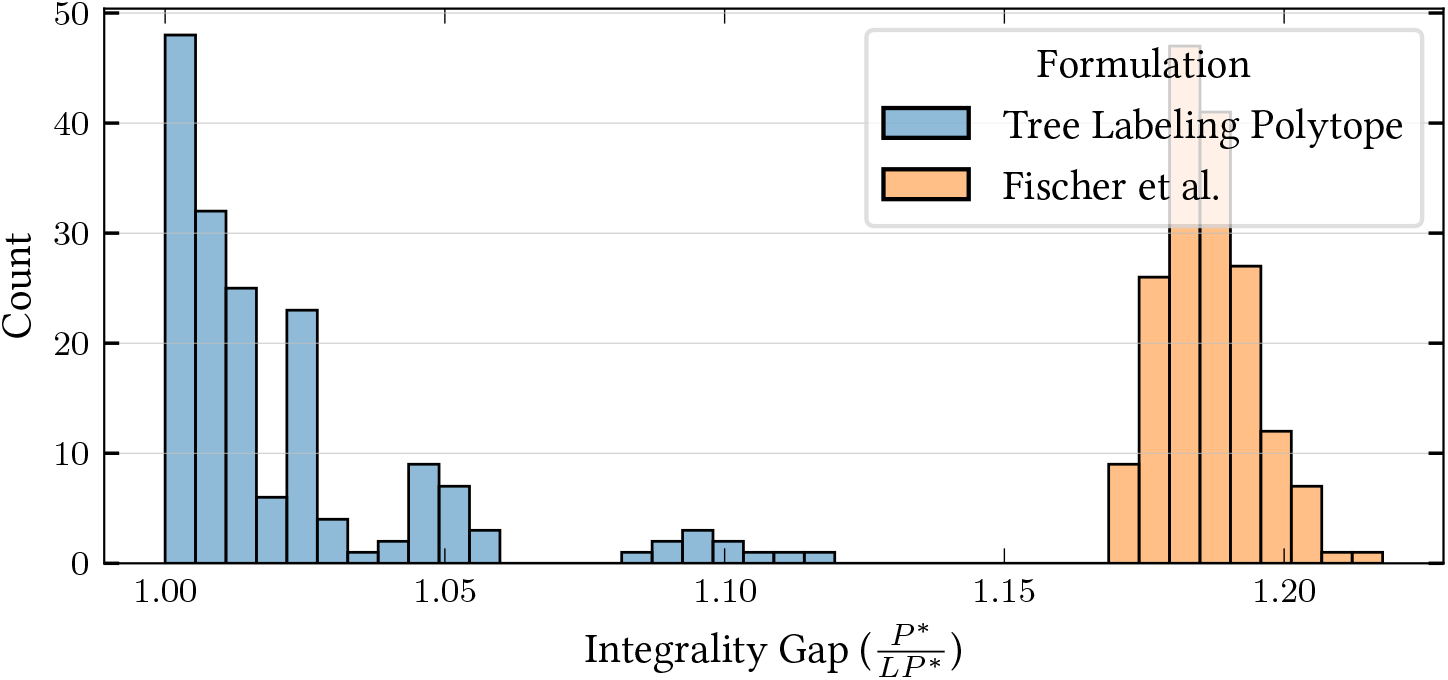
The integrality gap 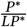 for the mixed integer programming formulations provided by the tree labeling polytope and Fischer et al. [31] across 198 simulated instances of the softwired small parsimony problem.

**Supplementary Figure 7:**
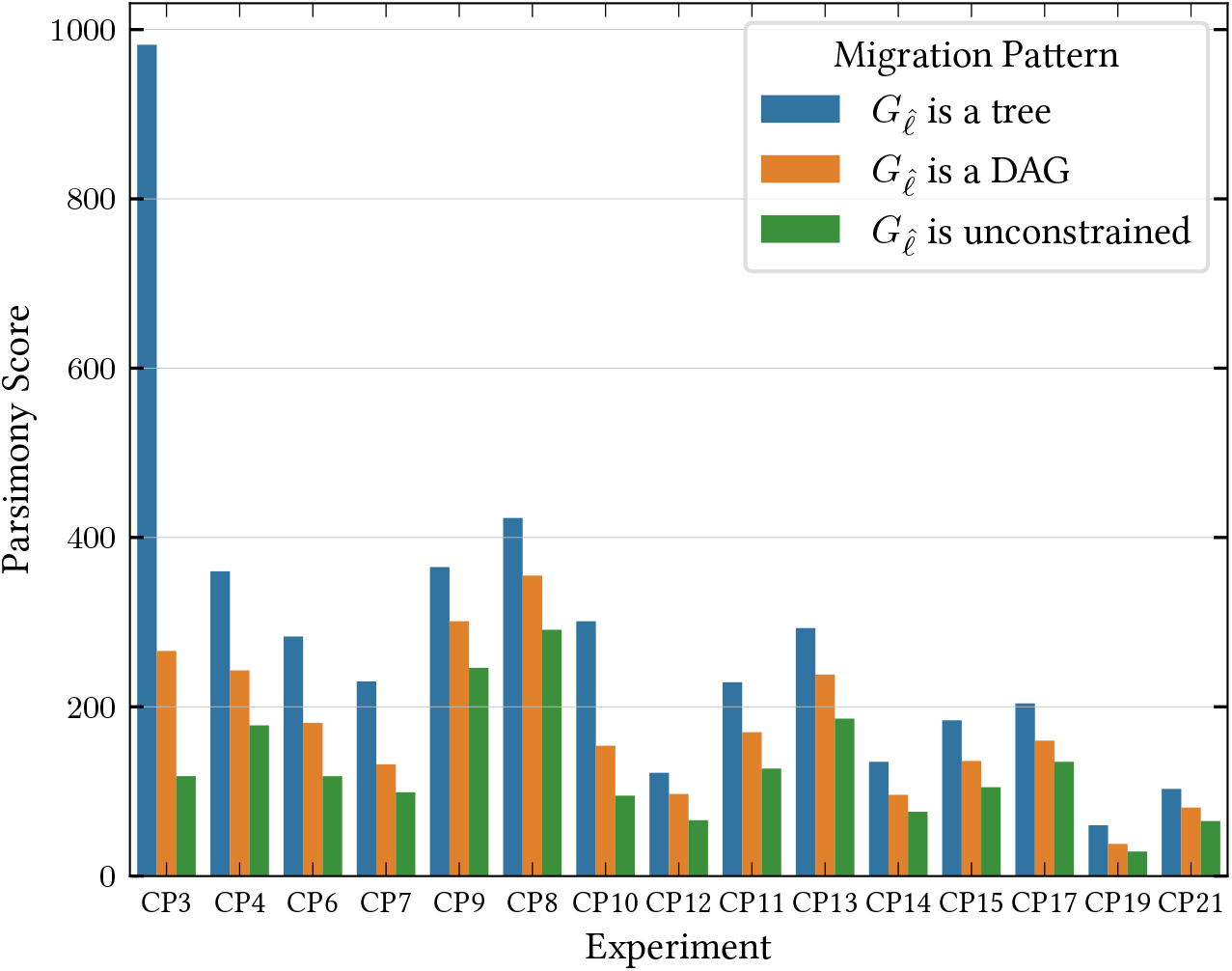
The parsimony scores *P*_0_, *P*_*D*_, *P*_*T*_ such that 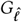 is unconstrained, a directed acyclic graph, and a tree, for the largest 15 clones in a mouse model of metastatic lung adenocarcinoma [55].

**Supplementary Figure 8:**
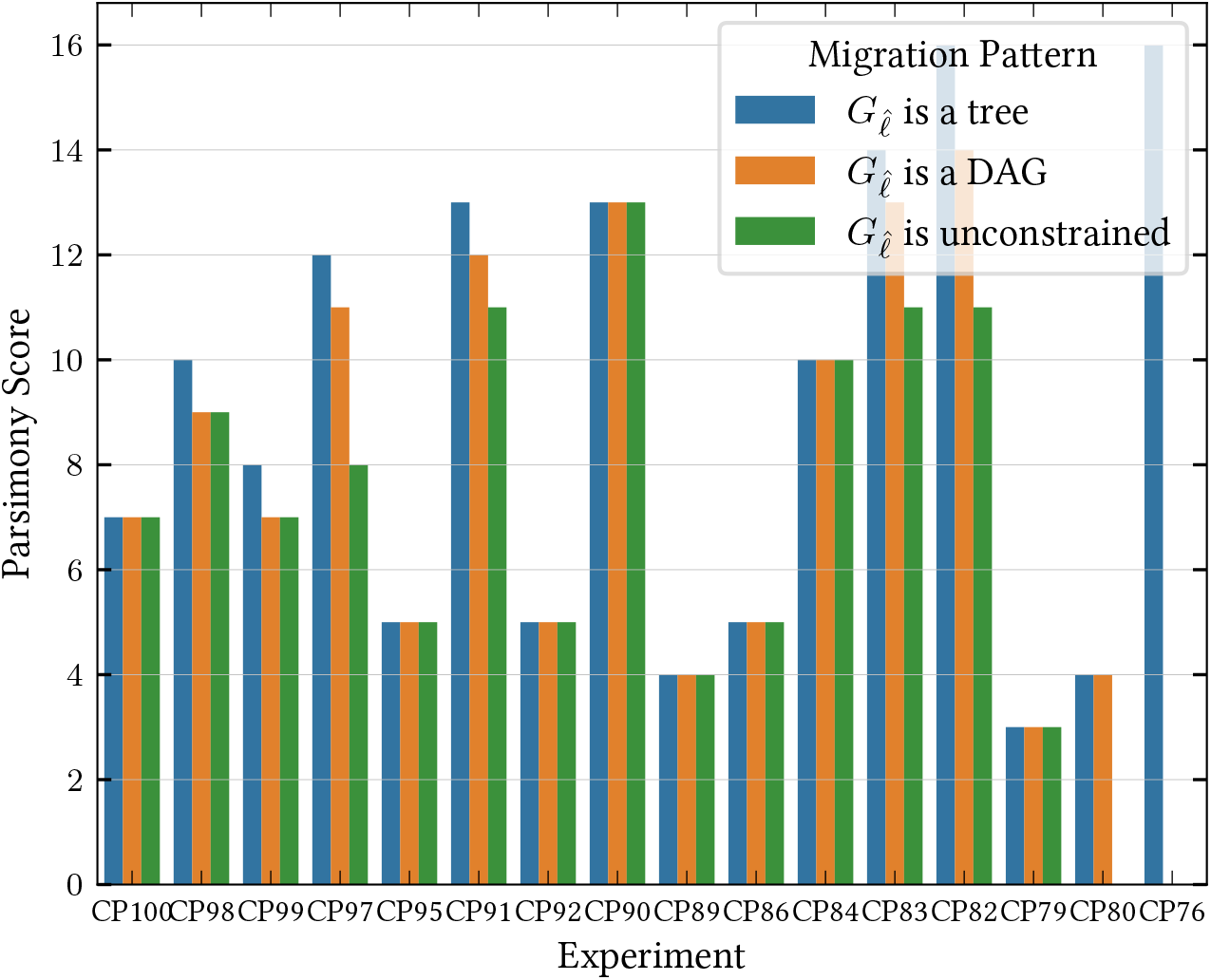
The parsimony scores *P*_0_, *P*_*D*_, *P*_*T*_ such that 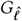 is unconstrained, a directed acyclic graph, and a tree, for the largest 15 clones in a mouse model of metastatic lung adenocarcinoma [55].

**Supplementary Figure 9:**
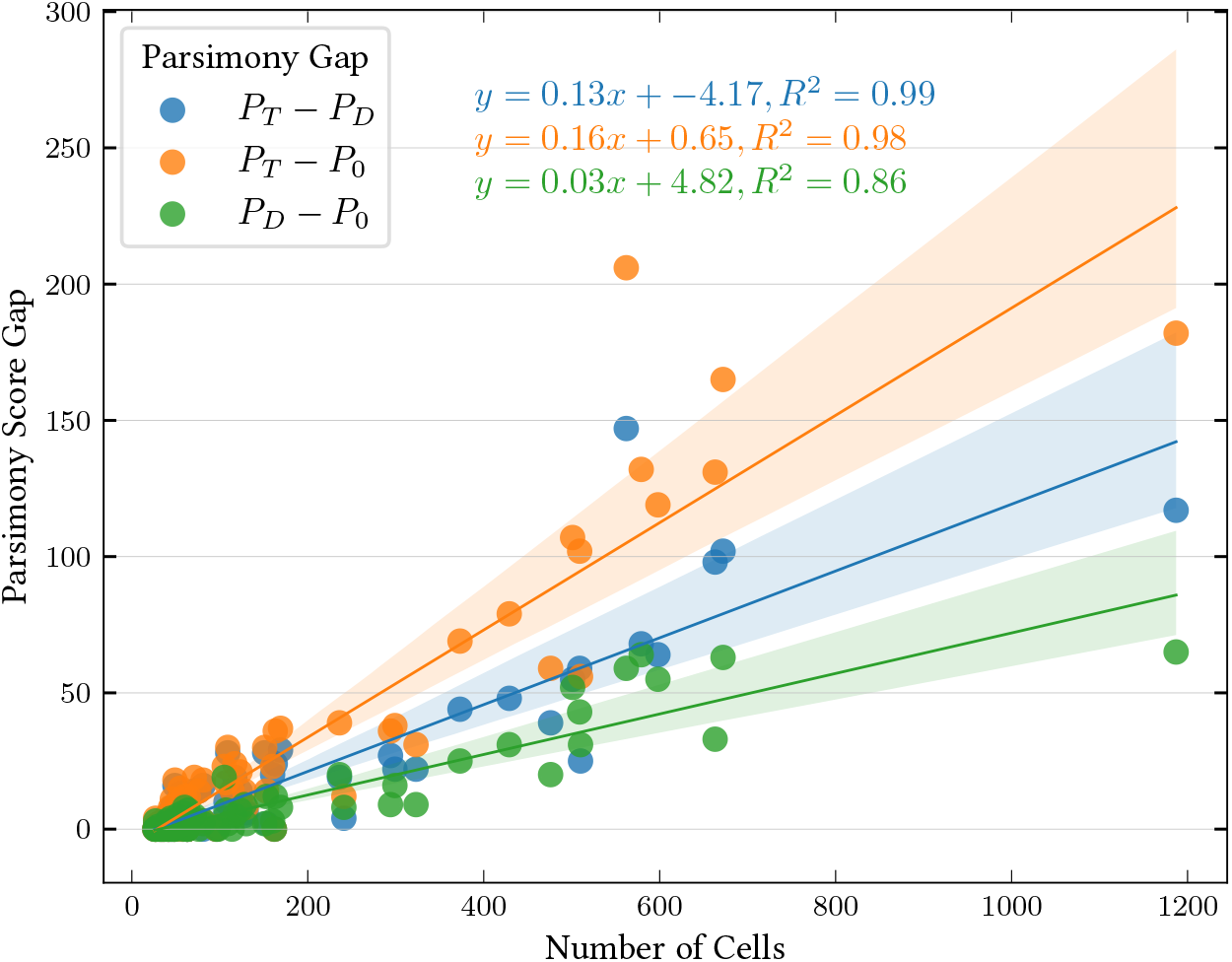
The parsimony gaps *P*_*T*_− *P*_*D*_, *P*_*T*_ −*P*_0_, and *P*_*D*_ −*P*_0_ versus the number of cells across all 83 clones in a mouse model of metastatic lung adenocarcinoma [55].

**Supplementary Figure 10:**
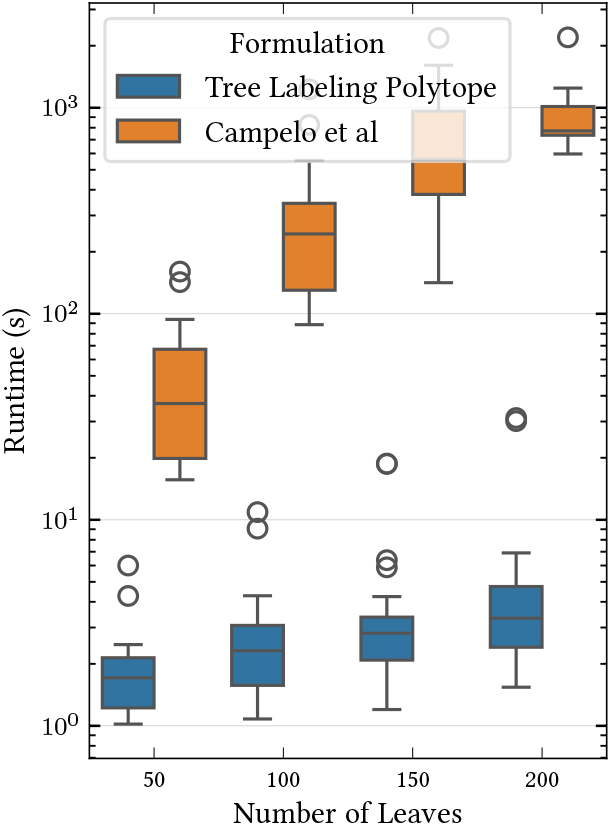
The wall-clock runtime (seconds) for the tree labeling polytope and Campelo [32] mixed integer linear programming formulations of the convex recoloring problem.

**Supplementary Figure 11:**
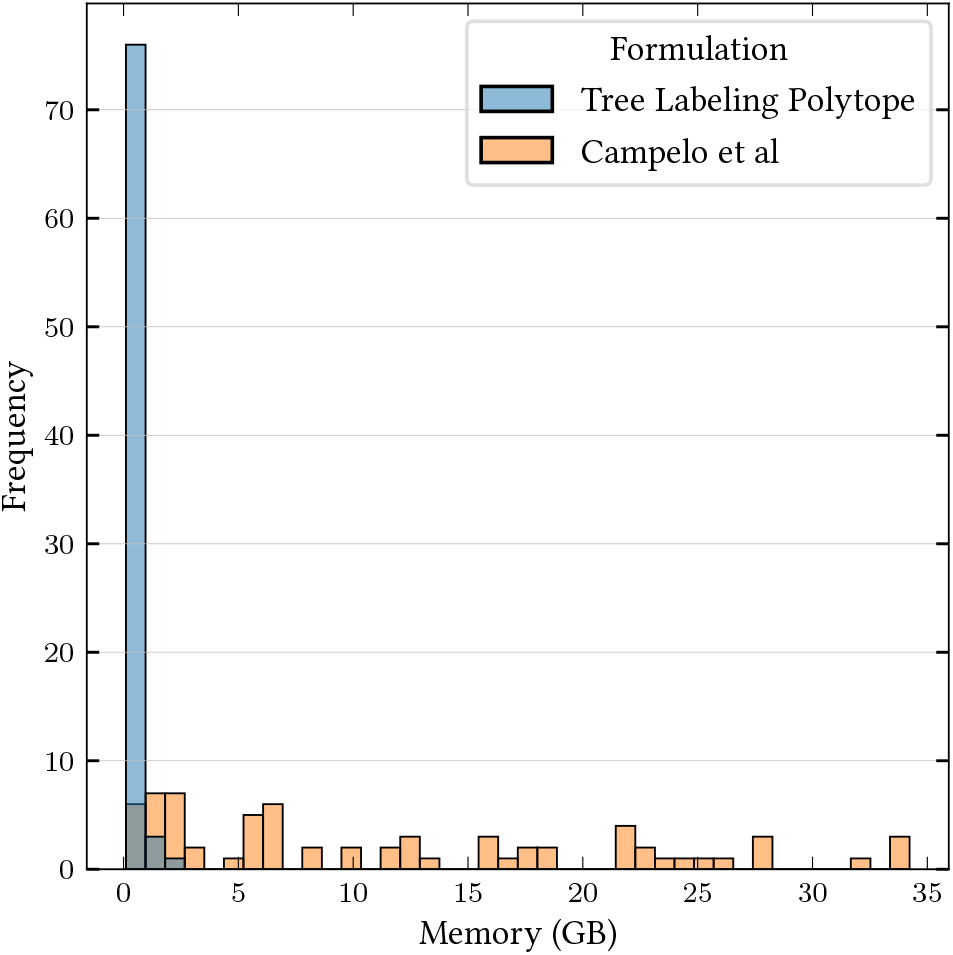
The memory usage in (GB) for the tree labeling polytope and Campelo [32] mixed integer linear programming formulations of the convex recoloring problem.

**Supplementary Figure 12:**
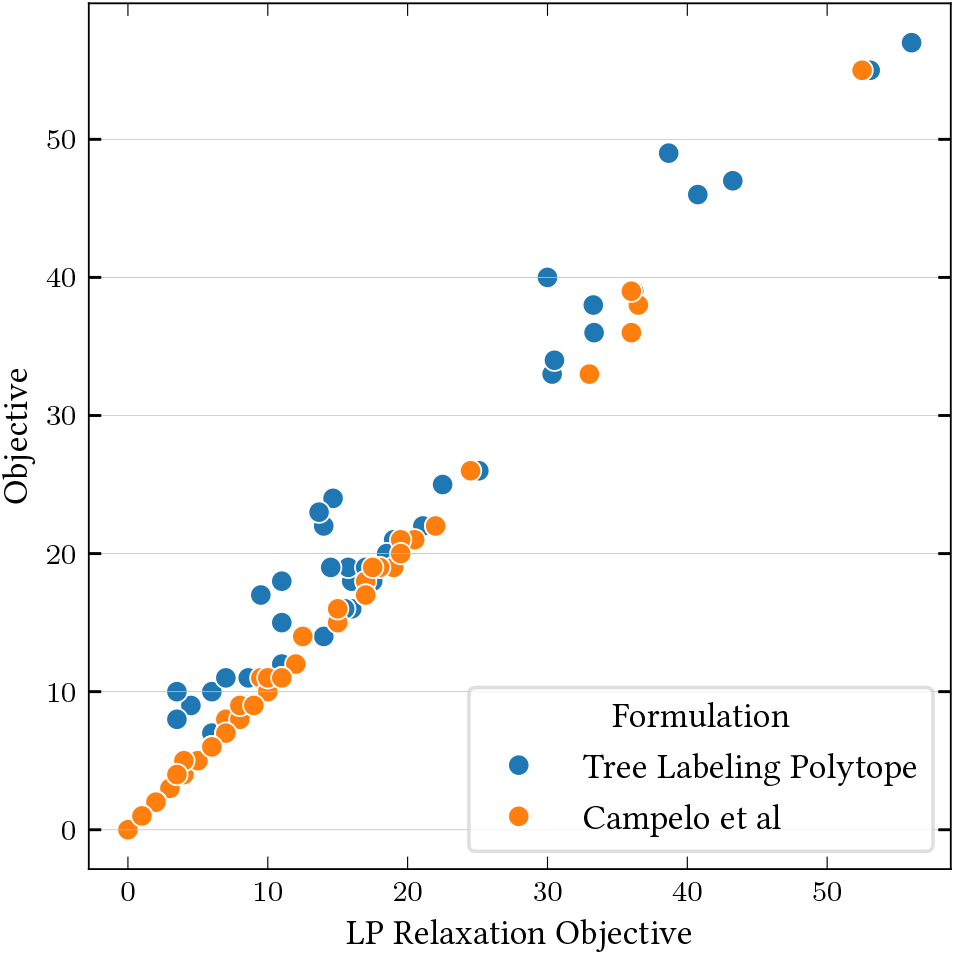
The linear programming relaxation objective versus the true objective value for the tree labeling polytope and Campelo [32] mixed integer linear programming formulations of the convex recoloring problem.

**Supplementary Figure 13:**
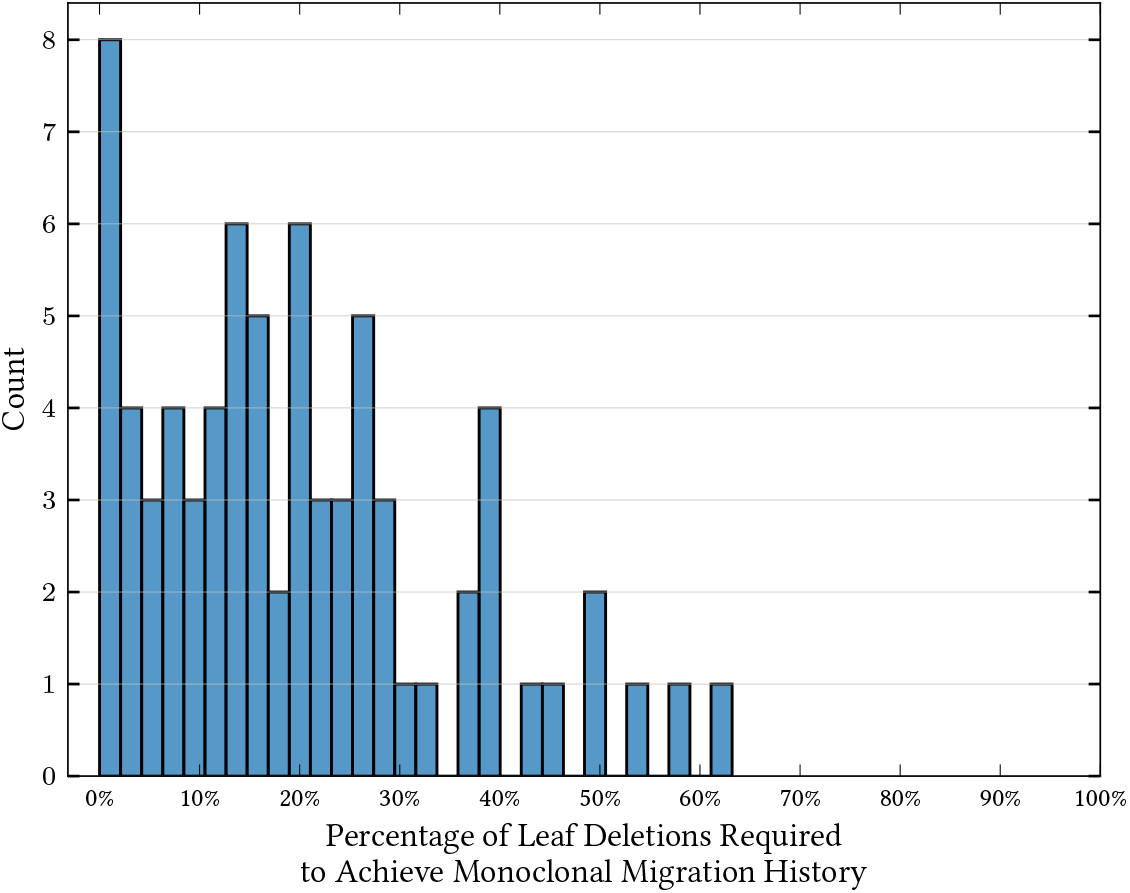
The percentage of cell deletions required to obtain a monoclonal explanation across the across all 83 clones in a mouse model of metastatic lung adenocarcinoma [55].

## Footnotes

1 A linear program min{*c*^*T*^ *x* : *Ax* ≤ *b*} which solves a combinatorial optimization problem *P* is called a *polyhedral description* for *P* since the feasible region *Q* = {*x* : *Ax* ≤ *b*} forms the convex polyhedron *Q* where each vertex of *Q* corresponds to a solution of *P*.

2 While this may appear to only model the case of a single phylogenetic character, typically, one assumes phylogenetic characters evolve independently.

3 A difference between our formulation of the parsimonious migration history problem and the formulation in MACHINA [11] is that we only consider migration graphs as opposed to *migration multigraphs*, which count the number of transitions between anatomical sites. We do this for simplicity of presentation, and our techniques apply to the case of migration multigraphs.

4 Though there is a simple set of linear inequalities describing the convex hull of these vectors.

5 A leaf labeling *ℓ* is *convex* if there exists an extension 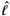 of *ℓ* such that for any two labels 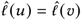, all vertices on the path *ν*_1_, …, *ν*_*k*_ from *u* to *ν* are labeled by 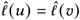.

